# Disordered but Different: Functional and Evolutionary Divergence of Transcription Factor Intrinsically Disordered Regions

**DOI:** 10.64898/2026.01.27.700206

**Authors:** Susie Song, Joshua M. Akey

## Abstract

Intrinsically disordered regions (IDRs) of proteins play key roles in multivalent interactions and the formation of biomolecular condensates. IDRs are widespread across the human proteome but are significantly enriched in transcription factor (TF) activation domains. However, it remains unclear why TF activation domains are enriched for IDRs and whether these regions are fundamentally distinct from IDRs in other proteins. Here, we comprehensively identify and analyze human IDRs and discover widespread functional, phenotypic, and evolutionary differences between TF and non-TF IDRs. Notably, in contrast to the broader proteome, TFs have evolved to become more disordered over time. Correspondingly, highly disordered TFs are more likely to regulate developmental processes, govern larger regulatory networks, and be subject to stronger regulatory constraints. TF IDRs are also enriched for pathogenic mutations relative to non-TF IDRs, and disorder content significantly predicts the mode of disease inheritance. Our results provide novel insights into how the evolution of gene regulation has uniquely shaped the molecular function and disease burden of TF IDRs.

## Introduction

Transcription factors (TFs) are modular proteins that make information encoded in the genome actionable by binding DNA and recruiting cofactors to activate or repress transcription^1^. TFs bind to regulatory elements, such as promoters and enhancers, through their structured DNA-binding domains and recruit cofactors through their intrinsically disordered activation domains. Unlike well-folded domains, intrinsically disordered regions (IDRs) of a protein lack a stable three-dimensional structure, and their biological functions are governed by the biochemical properties of their underlying amino acid sequences. Although TF activation domains are significantly enriched for IDRs^2,3^, disorder is a pervasive feature across the human proteome that contributes to diverse cellular processes, including transcription, signal transduction, and stress response^4–6^. A hallmark of IDRs is their ability to adopt dynamic conformational ensembles, allowing them to form transient, multivalent interactions^7–9^.

IDRs also promote liquid-liquid phase separation—a process that drives the formation of biomolecular condensates, which are membraneless compartments that regulate many biological processes^10–13^. Recent studies suggest that gene expression can be mediated by the assembly of transcriptional condensates, which are molecular hubs enriched for TFs and other components of the transcriptional machinery^14–16^. Experimental studies further implicate transcriptional condensates in enhancer activation, super-enhancer function, and the rapid transcriptional response to environmental perturbations^14,16–18^. Mechanistically, condensates have been proposed to act by locally enriching regulatory factors at target loci and altering TF binding kinetics, with downstream effects on transcriptional bursting dynamics^19,20^. Furthermore, aberrant formation of transcriptional condensates has been linked to neurodevelopmental disorders and cancer^21^.

Substantial progress has been made in delineating the functional roles that TF IDRs play in gene regulation and condensate dynamics^17,19^. Nonetheless, fundamental questions about the functional, phenotypic, and evolutionary significance of TF IDRs remain unresolved. In particular, TF activation domains are strikingly enriched for IDRs relative to the rest of the proteome^2,3^, but the origins of this enrichment are not well understood. Furthermore, although it has been shown that IDRs across the proteome often experience relaxed purifying and episodic positive selection^22–25^, few population genetics analyses have specifically focused on TF IDRs. Existing work suggests that extant patterns of genetic variation in TF IDRs are shaped by both positive and purifying selection^24–27^, but whether TF and non-TF IDRs differ in the tempo and mode of evolution remains unknown^28,29^.

Here, we address these gaps in knowledge by integrating proteome-wide structural annotations with large-scale population, evolutionary, and functional genomics datasets. We find pervasive differences in the characteristics of TF and non-TF IDRs, and how these differences impact the landscape of human disease. Collectively, these findings establish TF IDRs as a distinct and evolutionarily specialized component of the regulatory genome.

## Results

### Proteome-wide identification and analysis of IDRs reveals distinct properties of TF disorder

Transcriptional condensates are emerging as a key component of gene regulation^14–16^, and delineating the properties of TF IDRs is an essential step towards understanding how they form and regulate expression (Fig. 1a). To better understand the characteristics of TF IDRs and how they compare to disordered regions in other classes of proteins, we comprehensively identified IDRs in the human proteome (Fig. 1b). To this end, we downloaded the human proteome (*n* = 20,659 proteins) from UniProt^30^ and filtered out low-confidence entries (see Methods), resulting in a final set of 19,736 proteins. Of these, 8.2% (*n* = 1,612) were TFs^31^ (Fig. 1b).

**Fig. 1.**
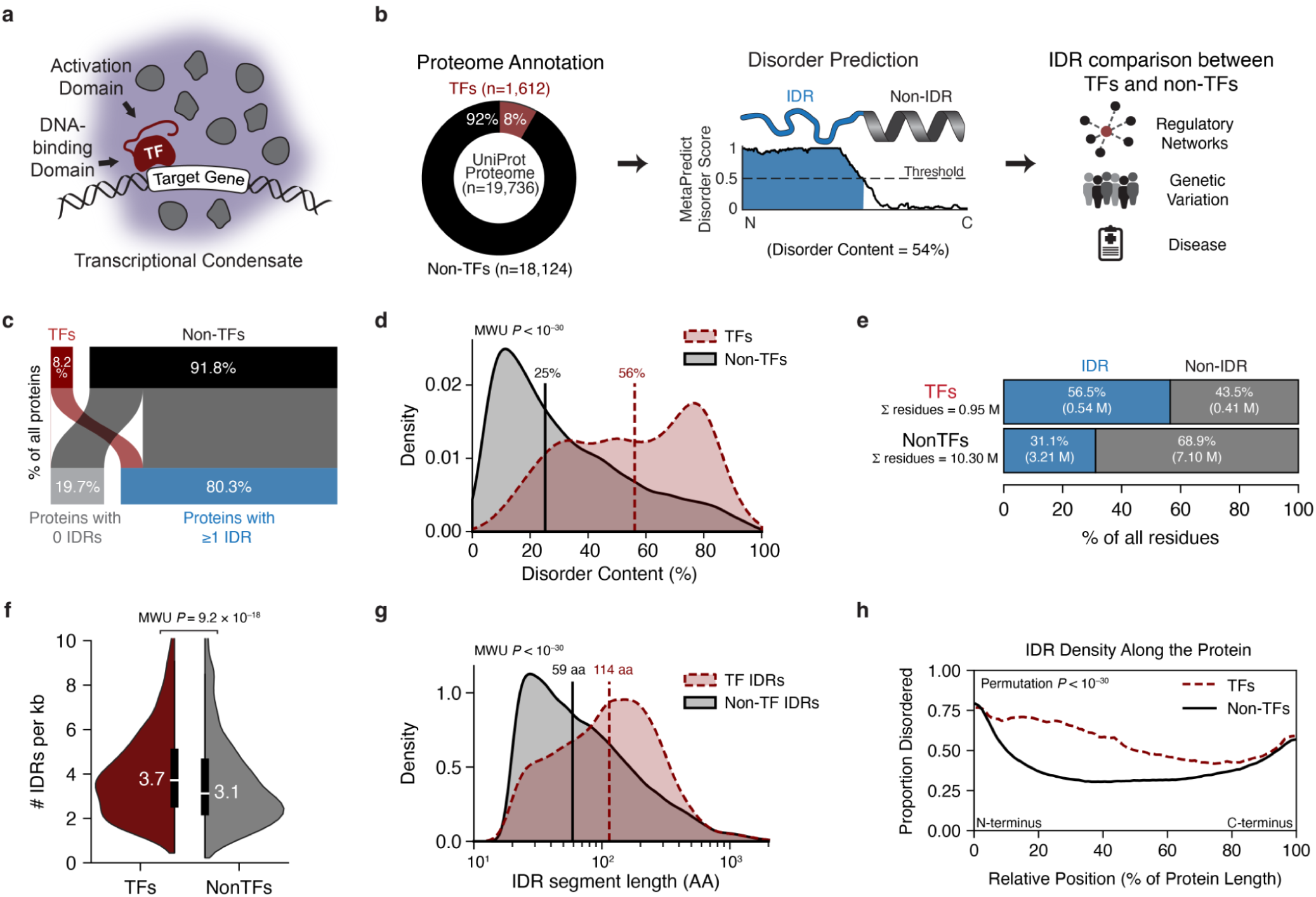
Identifying and characterizing TF and non-TF IDRs. **a**, Schematic of a TF with a structured DNA-binding domain (DBD) and disordered activation domain (AD) in a transcriptional condensate (purple). The disordered AD recruits multiple regulatory proteins (grey) to the target gene. **b**, Overview of the study workflow. UniProt human proteins were annotated as TFs or non-TFs. Residue-level disorder was predicted using MetaPredict, and IDR features were compared between TFs and non-TFs. **c,** Sankey diagram of the proportion of TFs (red) and non-TFs (black) with at least one predicted IDR (blue) versus no predicted IDRs (grey). **d**, Distribution of the per-protein disorder content among TFs (*n =* 1,581) and non-TFs (*n =* 13,180). Note, this analysis excludes proteins that are either completely ordered (IDR content=0%) or completely disordered (IDR content=100%). Vertical lines indicate group medians. The *P* value is from a two-sided Mann–Whitney U test. **e**, Fraction of all residues classified as an IDR or non-IDR, shown separately for TFs and non-TFs. M: million. **f**, Number of IDR segments per kb in TF and non-TF sequences. The median TF segment densities are labelled within the violin plots. The two-sided Mann–Whitney U test *P* value is shown. **g**, Length distribution of individual IDR segments in TFs (*n =* 3,248 IDRs) and non-TFs (*n =* 25,317 IDRs). The *P* value is from a two-sided Mann–Whitney U test. **h**, Position-dependent IDR density across normalized protein length, for TFs and non-TFs, showing the fraction of proteins disordered at each relative position. The *P* value reflects a permutation test of whether the disorder content between TFs and non-TFs across the normalized protein is different.

Next, we identified IDRs using MetaPredict^32,33^, a deep-learning-based consensus predictor shown to achieve high accuracy in independent benchmarking studies^34^. In total, we identified 24,638 IDRs (Fig. 1c), and 80.3% of all proteins (13,832 of 19,736) contained at least one IDR segment. Among proteins containing both ordered and disordered regions (Extended Data Fig. 1a), the median disorder content was significantly higher in TFs compared to non-TFs (56% vs 25%, respectively; Mann-Whitney U test, *P < 1*0^−30^; Fig. 1d). Across all TFs, 56.5% of amino acid residues fall within predicted IDRs, whereas for non-TF proteins, the corresponding value is 31.1% (Fig. 1e). Additionally, almost all TFs (99.3%) had at least one IDR segment (Extended Data Fig. 1b), and compared to non-TFs, they were significantly more likely to have two or more IDR segments, even after adjusting for gene length (Mann-Whitney U test, *P =* 9.2 × 10^−18^; Fig. 1f). We also confirmed this by stratifying proteins by length, and we observed a significant increase in IDR density across length quantiles for TFs compared to non-TFs (*P =* 2.1 × 10^−24^; Extended Data Fig. 1c). Additionally, the median TF IDR length was greater than that of non-TFs (Mann-Whitney U test, *P < 1*0^−30^; Fig. 1g). Finally, we examined the location of IDRs along the protein and found a significant bias towards the N- and C-termini for both TFs and non-TFs (permutation *P < 1*0^−30^; Fig. 1h). These data confirm previous inferences that TFs are enriched for disorder^2,3^, and additionally, demonstrate that IDRs occur more frequently and are longer in TFs compared to non-TFs.

To ensure that our inferences are robust to the choice of IDR prediction method, we also identified disordered regions with AIUPred^35^, which relies on pairwise contact energy estimations and has been shown to perform well in benchmarking studies^34^ (see Methods). IDR predictions were highly concordant between MetaPredict and AIUPred (Extended Data Fig. 1d-f). For example, the number of amino acids predicted to be disordered per protein and the proportion of a protein predicted to be disordered were significantly correlated between the two methods (Pearson *r* = 0.94 and 0.78, respectively; *P < 1*0^−30^ for both correlations; Extended Data Fig. 1d-e). Similarly, the Hamming distance, which quantifies the fraction of residues with discordant predictions of disorder (see Methods), was small (median = 0.04; Extended Data Fig. 1f). Perhaps most importantly, all of the analyses described below resulted in the same qualitative and quantitative inferences when using either the MetaPredict or AIUPred callsets (Extended Data Fig. 1g-h). Thus, these data demonstrate that our results are robust to the specific method used to identify IDRs.

### TFs evolved to become more disordered

The disorder-to-order model of protein evolution posits that newly emerged proteins are initially disordered and progressively acquire folded domains with time^36–38^ (Fig. 2a). To assess whether this paradigm applies to TFs, we tested whether there was a relationship between the disorder content of TFs and non-TFs as a function of gene age. We obtained estimates of gene age from GenOrigin^39^, which dates protein-coding genes across 565 species using Ensembl/Ensembl Genomes gene families and parsimony-based inference on a species timetree. Under the disorder-to-order model, we would expect the oldest proteins to have the least disorder and the youngest proteins to have the most. The distribution of disorder content for non-TFs decreases as gene age increases (Kruskal-Wallis, *P < 1*0^−30^; Fig. 2b), consistent with the disorder-to-order model. In contrast, TFs follow the opposite pattern, with disorder content increasing with gene age (Kruskal-Wallis *P < 1*0^−30^; Fig. 2b). Thus, the oldest TFs (>1000 Mya) are enriched for disorder, whereas the oldest non-TFs are depleted of disorder. These data suggest that TF IDRs have experienced a unique evolutionary trajectory compared to non-TF IDRs, in which disorder is not lost but instead maintained or expanded over evolutionary time (Fig. 2c).

**Fig. 2.**
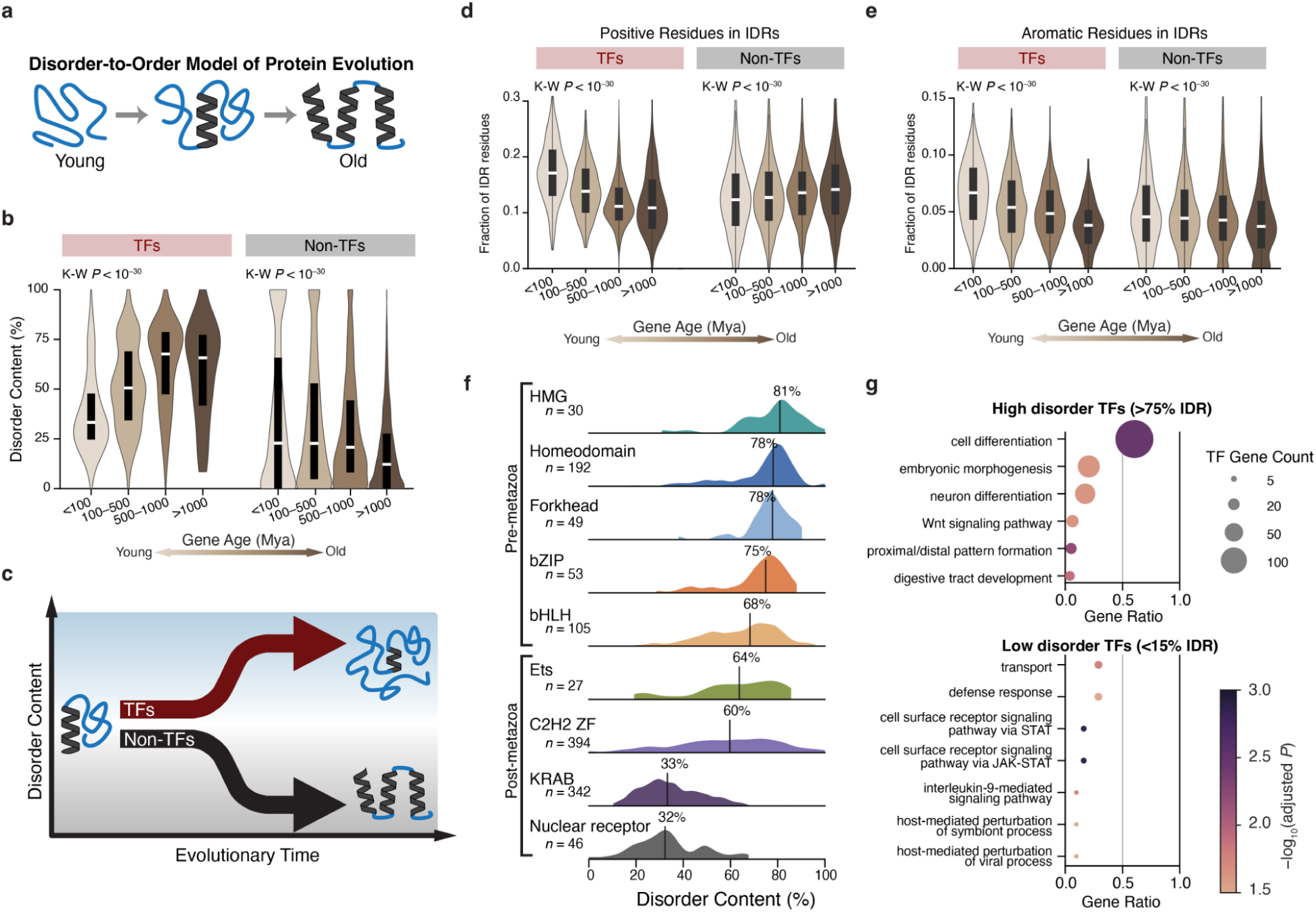
Evolutionarily ancient TFs are more disordered. **a,** The “disorder first” hypothesis of protein evolution, in which a peptide begins disordered (blue) and acquires secondary structure (grey) over time. **b,** Disorder content stratified by inferred gene age for TFs and non-TFs. Evolutionarily ancient TFs are enriched for disorder, whereas the opposite trend is seen in non-TFs. *P* values are from the Kruskal–Wallis test within each protein group. White dots denote medians, and lines denote the interquartile range (IQR). **c,** Summary schematic of the observed divergence between TFs and non-TFs: TF disorder content is maintained or ever expanded over time, whereas non-TFs show the opposite trend, consistent with distinct selective pressures on IDRs in regulatory versus non-regulatory proteins. **d–e,** Composition of IDRs across gene ages, showing that the fraction of (**d**) positively charged residues (K, R, H) and (**e**) aromatic residues (F, W, Y) within IDRs varies with age in both TFs and non-TFs (Kruskal–Wallis tests), suggesting that not only the amount of disorder but also the biochemical characteristics of IDRs shifts across evolutionary time. **f,** Disorder content split by the largest TF DNA-binding domain families, organized into pre- and post-metazoan families. Vertical lines denote medians. **g,** GO Biological Processes enrichment using a subset of GO terms (levels 3-5 terms) in TFs with high disorder (IDR content>75%; *n =* 351; top) and low disorder (IDR content<15%; *n =* 31; bottom). Dot plots show enriched GO terms (*y* axis). Gene ratio (*x* axis) is the proportion of TFs in the queried set annotated to a given term. The full TF list was used as the background gene set (*n =* 1,008). Dot size indicates the number of TFs annotated to each term, and dot color indicates *−log_10_*(Benjamini–Hochberg adjusted *P* value).

Despite the striking differences in IDR content and gene age between TFs and non-TFs, several factors can confound estimates of gene age^38,40–42^. Therefore, we performed additional analyses to evaluate the robustness of these patterns. First, we repeated our analysis of disorder content across age strata using an independent catalog of gene ages^43^, which corrects for biases caused by failures in homology detection (Extended Data Fig. 2). Across phylostrata, we observed consistent trends for both TFs and non-TFs (Extended Data Fig. 2a). TFs that arose in Gnathostomata or earlier consistently exhibited higher disorder content than younger TFs (Mann-Whitney U test, *P =* 1.5×10^−10^; Extended Data Fig. 2b), whereas non-TFs showed the opposite pattern (Mann-Whitney U test, *P < 1*0^−30^; Extended Data Fig. 2b). Second, these disorder-age relationships were unchanged when stratifying the GenOrigin-based gene age estimates by sequence conservation, protein length, GC content, expression level, and tissue specificity (Extended Data Fig. 2c–g), indicating that these potential confounding factors do not explain the elevated disorder in ancient TFs. Third, we fit a multivariable linear model (OLS) of gene age on disorder content, TF status, and their interaction, adjusting for protein length, GC content, mean expression, and tissue specificity (see Methods). The disorder-by-TF interaction was strongly positive *β* = 12.97, *P < 1*0^−30^) indicating that the age–disorder slope differs between TFs and non-TFs after covariate adjustment. In this model, disorder content was negatively associated with gene age among non-TFs (*β* = −5.48, 95% CI −6.12 to −4.85), consistent with the canonical disorder-to-order trend, whereas TFs showed a positive association (*β* = 7.49, 95% CI 5.60 to 9.39). Collectively, these results support a model in which TFs do not follow the canonical disorder-to-order trajectory and instead maintain, or expand, disorder content over evolutionary time.

We next investigated factors that may have contributed to the increasing disorder content over time in TFs. Specifically, we examined how the amino acid composition of IDRs changes over time, as such shifts can reveal selective pressures acting on IDR biochemistry. We tested five categories of amino acid residues: polar, aromatic, aliphatic, positively charged, and negatively charged. Of these, we found significant trends for positively-charged (Fig. 2d) and aromatic residues (Fig. 2e). The youngest TF IDRs are significantly enriched for positively charged residues (lysine, arginine, histidine; Kruskal-Wallis, *P <* 10^−30^; Fig. 2d) and aromatic residues (phenylalanine, tyrosine, tryptophan; Kruskal-Wallis, *P <* 10^−30^; Fig. 2e), consistent with the hypothesis that aromatic residues arose later in the evolution of the genetic code^44–46^. Older TF IDRs, in contrast, are relatively depleted of these residues (Fig. 2d-e). Non-TF IDRs show the opposite charge trajectory—an increase in positive residues with age—and only modest changes in aromatic content (Fig. 2d-e). The distinct charge trajectories imply that the physicochemical pressures shaping IDRs have diverged between TFs and non-TFs. In summary, these data show that TFs do not follow the classic disorder-to-order model of protein evolution^36–38^, and that the biochemical grammar of their IDRs has itself evolved, potentially reflecting changing requirements for regulatory complexity and interaction capacity.

### TF Families Exhibit Marked Heterogeneity in Disorder

TFs are organized into families based on their DNA-binding domain^31,47^, providing a natural framework to compare disorder content across functionally distinct TFs. We observed that TF families have marked heterogeneity in levels of disorder (Fig. 2f). Evolutionarily ancient families, such as high mobility group (HMG) and homeodomain TFs, had the highest levels of disorder (median percent of disordered residues per protein of 81% and 78%, respectively; Fig. 2f). In contrast, the nuclear receptor and KRAB family of TFs exhibit the lowest amount of disorder (median percent of disordered residues per protein of 32% and 33%, respectively; Fig. 2f). Additionally, there is variation in IDR length, number, and magnitude of terminal bias among TF families (Extended Data Fig. 3a-e).

We next investigated whether TF disorder content was associated with specific biological processes. For example, many developmental TFs, such as HOXA2 (86%), HOXA3 (88%), FOXO1 (90%), and FOXO3 (90%), are highly disordered (defined as ≥75% of amino acid residues predicted to be disordered across the protein). Indeed, GO enrichment analysis revealed that highly disordered TFs are significantly enriched (FDR ≤ 5%) for GO terms related to development and metabolism (Fig. 2g; Extended Data Fig. 4a). As expected, the average amount of disorder for TFs annotated with the GO term “developmental process” (GO:0032502) is significantly higher than the rest of the TFs (Mann-Whitney U-test, *P < 1*0^−30^; Extended Data Fig. 4b). Non-TFs annotated as developmental also have a significantly higher average disorder content compared to other non-TF proteins (Mann-Whitney U-test, *P =* 3.4×10^−9^; Extended Data Fig. 4b), but the magnitude of the difference was much smaller compared to TFs. Furthermore, pioneer TFs^48–50^, which initiate transcription by binding to nucleosomes and promoting chromatin accessibility, are significantly enriched for disorder relative to other TFs (Mann-Whitney U-test, *P =* 2.7×10^−4^; Extended Data Fig. 4c). Overall, these data suggest that highly disordered TFs contribute to core biological processes (Fig. 2g, Extended Data Fig. 4a). In contrast, ordered TFs (≤15% of the protein is disordered) tend to be enriched for signaling-related terms (Fig. 2g). Examples of highly ordered TFs include the STAT family of signaling TFs (*STAT1, STAT3, STAT4*) and the IRF family of immune related TFs (*IRF4, IRF8*).

### The regulatory characteristics of TFs are shaped by their disorder content

To evaluate whether the disorder content of TFs is associated with functional characteristics, we examined protein-protein interaction (PPI) networks and gene regulatory networks (GRNs). We hypothesized that highly disordered TFs would interact with a larger protein network. Using the STRING PPI database, we found a significant positive correlation between TF disorder content and the number of proteins in its functional association network (indirect interactions) (Spearman’s *ρ =* 0.36, *P <* 10^−30^; Fig. 3a). In contrast, non-TFs exhibit a significantly negative correlation between disorder content and number of proteins in its functional association network (Spearman’s *ρ = −*0.14, *P <* 10^−30^; Fig. 3a). This finding is consistent with the hypothesis that TF activation domains facilitate multiple interactions, enabling flexible coordination of transcriptional cofactors and machinery.

**Fig. 3.**
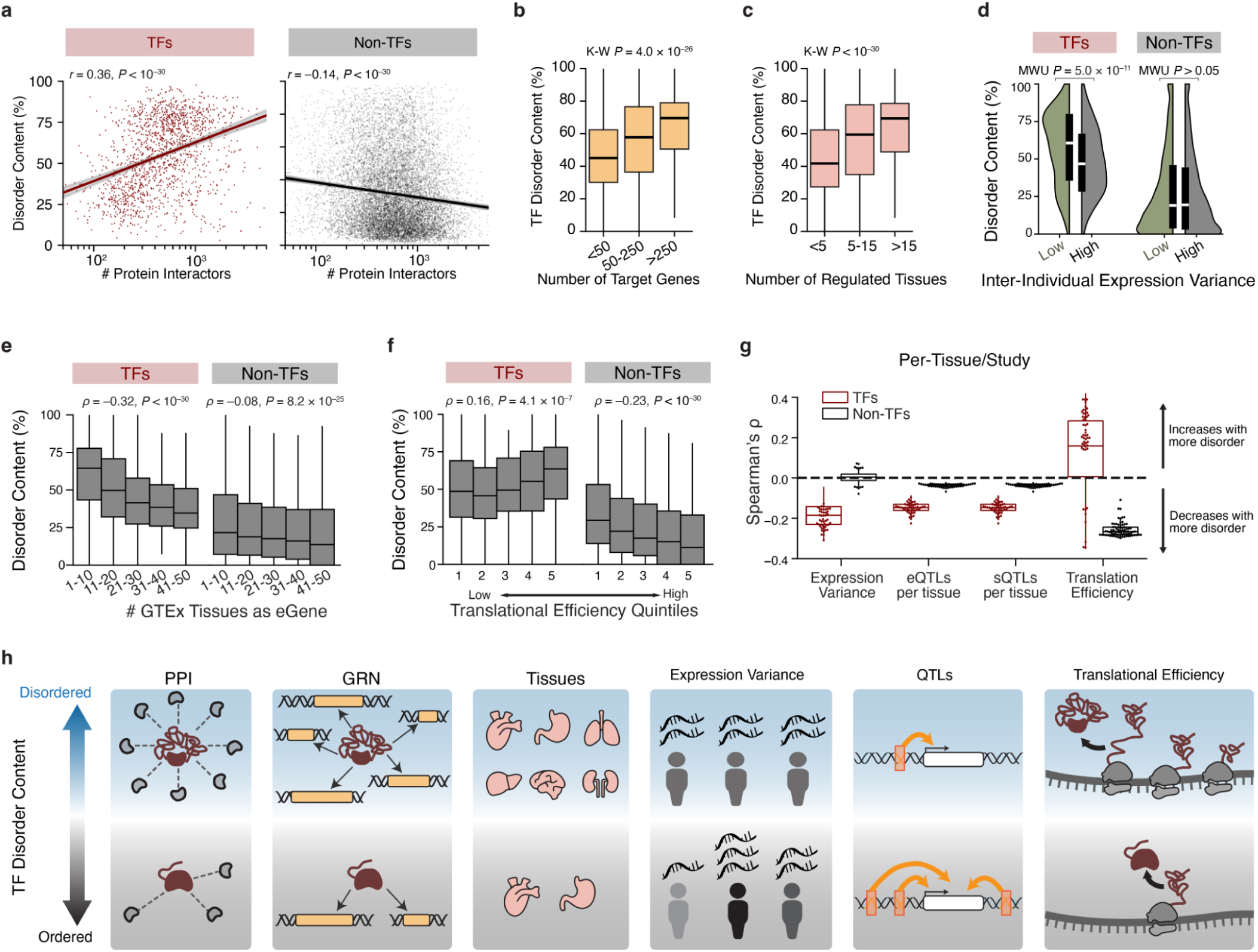
Disorder content of TFs is associated with functional genomics phenotypes. **a,** Disorder content as a function of protein–protein interaction (PPI) degree for TFs and non-TFs. Interactions include direct interactions and indirect associations from STRING^77^. TF disorder content increases with the number of interactors in its functional association network, whereas non-TFs show a weak negative association. Lines indicate linear fits with 95% confidence bands. **b,** TF disorder content increases with the number of target genes assigned to each TF in GRNs. The Kruskal–Wallis test *P* value compares bins of target gene size. Per box TF counts from left to right are *n =* 420, 384, 255. **c,** TF disorder content also increases with the number of tissues in which a TF is inferred to be an active regulator (controlling a GRN regulon). The Kruskal–Wallis test *P* value compares bins of the number of regulated GTEx tissues. Per box TF counts from left to right are *n =* 655, 484, 356. **d,** Disorder content stratified by inter-individual expression variance in GTEx^78^ for TFs (high-variance TFs *n =* 510; low-variance TFs *n =* 421) and non-TFs (high-variance non-TFs *n =* 5,270; low-variance non-TFs *n =* 5,194); high-variance TFs exhibit increased disorder content, whereas non-TFs show no significant difference (two-sided Mann–Whitney U tests). **e**, Disorder content as a function of the number of GTEx tissues in which a gene is detected as an eGene. Disorder content decreases with increasing eGene breadth for both TFs and non-TFs. Per box sample sizes for TFs from left to right are *n =* 879, 239, 132, 110, and 101; non-TFs: *n =* 7,537, 3,736, 2,143, 1,404, and 1,225. **f**, Disorder content across translational efficiency^56^ quintiles; TF disorder content increases with translational efficiency, whereas non-TFs show the opposite trend. *n =* 190 per TF quintile and *n =* 1,849 per non-TF quintile. **g**, Per-study association summary. For each study/tissue, Spearman correlations were computed between disorder content and each metric (expression variance, eQTLs per tissue, sQTLs per tissue, and translational efficiency); boxplots summarize the distribution of per-tissue correlation coefficients for TFs and non-TFs, with the dashed line indicating no association. The sample sizes (number of tissues or studies) for each group, from left to right, are *n =* 57, 50, 50, and 78. **h,** Conceptual summary: higher TF disorder content is associated with larger protein functional association networks, broader regulatory targeting within GRNs, more tissues regulated, less expression variance, fewer e/sQTLs and greater translational efficiency.

We performed several additional analyses to evaluate the robustness of the positive relationship between disorder content and PPI connectivity for TFs (Extended Data Fig. 5a-d). For example, when quantified as the total number of disordered amino acids rather than as a percentage, TF disorder content remained significantly correlated with the number of PPIs (Spearman’s *ρ =* 0.22, *P =* 2.5×10^−19^; Extended Data Fig. 5a). We also found that the relationship between disorder content and PPIs for TFs and non-TFs was robust to the confidence threshold used to define an interaction (see Methods), for both functional network interactions and when restricted to physical interactions (Extended Data Fig. 5b). Furthermore, we repeated the analysis with two additional PPI databases: the HPA^51^ and BioGRID^52^, and observed the same trends (Extended Data Fig. 5c-d). Thus, these results suggest that the relationship between disorder and the number of interacting proteins is not an artifact of database coverage or interaction type.

Next, we examined GRNs inferred from the GTEx Project^53^ to determine whether there is a relationship between TF disorder content and the size of the GRN it regulates^54^ (see Methods). We observed a significant positive correlation between TF disorder content and the total number of target genes in GRNs across all tissues it regulates (Spearman’s *ρ =* 0.28, *P =* 6.6×10^−24^; Fig. 3b), across a broad range of GTEx tissues (Spearman’s *ρ =* 0.32, *P <* 10^−30^; Fig. 3c). We also confirmed that these correlations were also observed in GRNs inferred from an independent dataset^55^ (Extended Data Fig. 5e-f).

### Highly disordered TFs are under stronger regulatory constraint

We next explored whether TF disorder content was associated with characteristics of its expression. TFs expressed in fewer tissues (more narrow expression breadth, as annotated by HPA^51^) are enriched for disorder content (Kruskal-Wallis, *P <* 10^−30^; Extended Data Fig. 6a). Highly disordered TFs are more likely to exhibit tissue-specific expression (uneven expression across tissues) in bulk RNA^51^ (Mann-Whitney U test, *P <* 10^−30^; Extended Data Fig. 6b), whereas non-TFs show no such relationship (Mann-Whitney U test, *P* > 0.05; Extended Data Fig. 6b). Highly disordered TFs are also enriched for uneven expression across cell types, or cell type-specific expression^51^ (Mann-Whitney U test, *P =* 5.1×10^−5^; Extended Data Fig. 6c). Additionally, we found that highly disordered TFs exhibit significantly less gene expression variation across individuals (Mann-Whitney U test, *P =* 5.0×10^−11^, Fig. 3d), suggesting that the transcription of these genes is tightly regulated.

Since purifying selection purges genetic variation that has deleterious effects on gene expression, we hypothesized that highly disordered TFs would have fewer expression quantitative trait loci (eQTLs). To test this hypothesis, we examined the fine-mapped eQTL sets from GTEx^53^ and found that TF eGenes (i.e., TFs whose transcript abundance is influenced by one or more eQTL) that are present in many tissues are depleted of disorder (Fig. 3e). Additionally, genes regulated by many independent credible sets, indicating multiple causal genetic variants, are depleted for disorder (Extended Data Fig. 6d). Thus, highly disordered TFs are depleted of eQTL. Interestingly, the same trend was also observed with fine-mapped TF splicing QTLs (sQTLs) (Extended Data Fig. 6e-f), suggesting that both gene expression and isoform selection are tightly regulated in highly disordered TFs, and that inter-individual variation of these molecular phenotypes is poorly tolerated.

To further investigate the stronger regulatory constraints of highly disordered TFs, we analyzed patterns of translational efficiency^56^ (see Methods). We found a significant positive correlation between translational efficiency of TF mRNAs and TF disorder content (Spearman’s *ρ* = 0.16, *P =* 4.1×10^−7^, Fig. 3f). In contrast, non-TFs showed the opposite pattern, with highly disordered proteins having lower levels of translational efficiency (Spearman’s *ρ* = −0.23, *P* < 10^−30^, Fig. 3f). Importantly, correlations between disorder content and expression variation across individuals, number of eQTL and sQTL per tissue, and translational efficiency were largely stable across tissues (Fig. 3g). Together, these data indicate that highly disordered TFs are subject to stronger regulatory constraints, and because of their important biological functions have evolved robust regulatory architectures to buffer the effects of genetic variation or transcriptional noise (Fig. 3h).

In summary, the degree of intrinsic disorder in TFs is strongly associated with their functional characteristics. TFs with higher levels of disorder tend to participate in more protein–protein interactions, regulate larger gene regulatory networks, and are regulators across a broader range of tissues (Fig. 3h). These findings are also consistent with the observation that ancient TFs are more highly disordered and that intrinsic disorder contributes to their broad regulatory roles.

### Population genetics characteristics of TF IDRs

We next asked whether the different characteristics of TF and non-TF IDRs described above are also reflected in patterns of human genomic variation. To this end, we leveraged the gnomAD^57^ database to compare characteristics of genetic variation in ordered and disordered protein domains between TFs and non-TFs. The gnomAD database is a powerful resource for population-genetic inference, as it aggregates sequence data from up to 1 million individuals.

For both TFs and non-TFs, SNVs and indels are significantly enriched in IDRs relative to non-IDRs (Binomial exact test, *P < 1*0^−30^; Fig. 4a). Specifically, in TFs, 60% of SNVs and 62% of indels occur in IDRs, significantly exceeding the 57% of disordered TF coding sequence (Fig. 4a). For TFs, these numbers translate to a variant density of 355.1 SNVs/kb in IDRs and 307.2 SNVs/kb in non-IDRs. Non-TFs exhibit a similar enrichment, with 32% of SNVs and 36% of indels falling in IDRs, despite IDRs constituting only 31% of their coding genome (Binomial exact test, *P < 1*0^−30^; Fig. 4a). For non-TFs, these numbers translate to a variant density of 339.0 SNVs/kb in IDRs and 322.8 SNVs/kb in non-IDRs. Thus, as expected, disordered regions of both TFs and non-TFs are more tolerant of mutations compared to ordered regions of the protein. Furthermore, although ordered regions of TFs are subject to higher overall levels of purifying selection than non-TFs, TF IDRs are less constrained than IDRs of non-TFs.

**Fig. 4.**
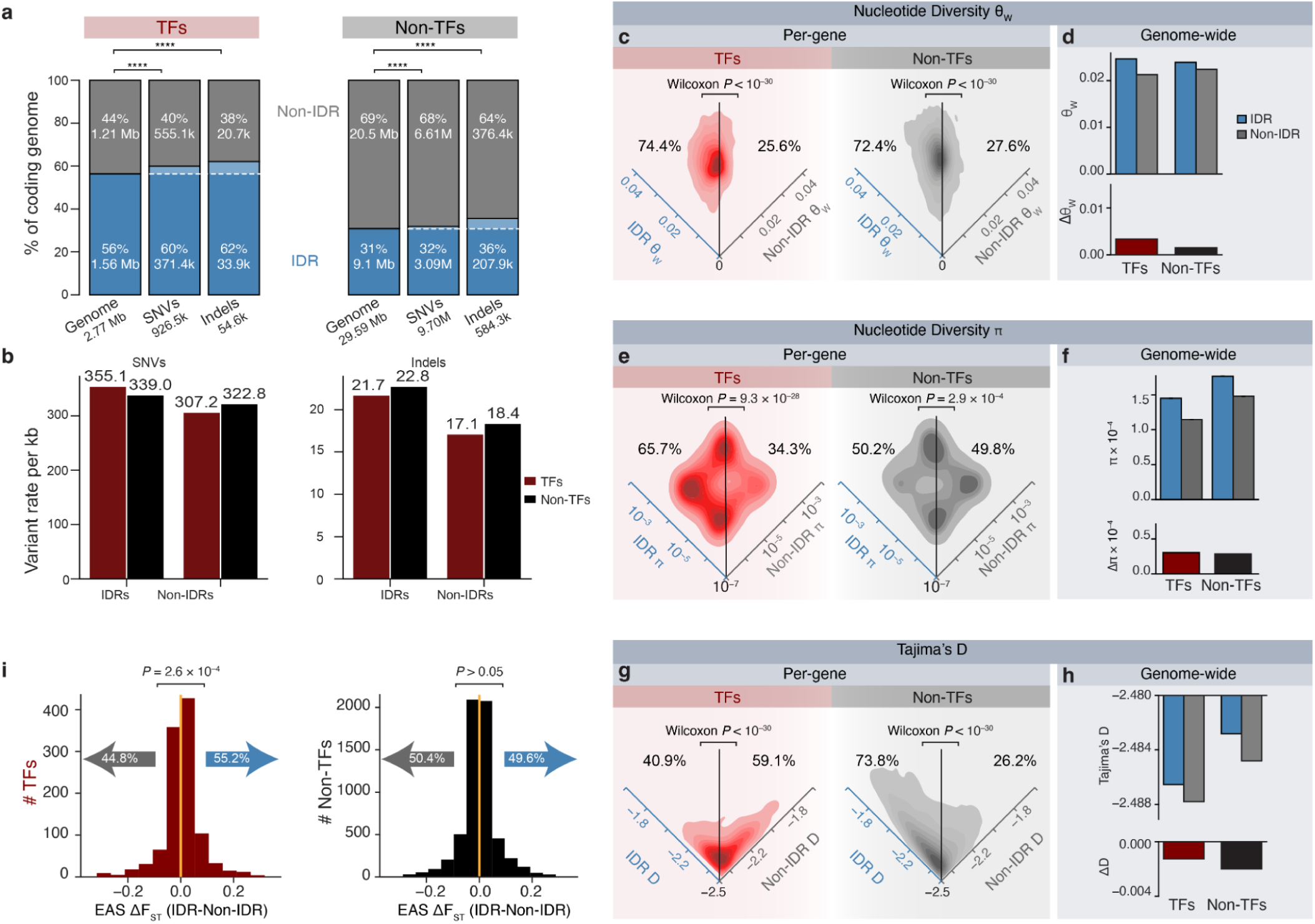
Population genetics of TF and non-TF IDRs. **a,** TF IDRs are enriched for SNVs and Indels in gnomAD v4.1. Total genomic region, SNV, and indel counts are shown, split between TF Non-IDRs (grey) and IDRs (blue). *P* values were obtained from the binomial exact test. **** indicates *P < 1*0^−30^. **b**, SNV and indel rates. kb: kilobase. **c,** Nucleotide diversity θ_W_, in TFs (*n =* 1,483; left) and non-TFs (*n =* 10,182; right) with between 10% and 90% disorder content. θ_W_ was calculated separately for the protein’s IDR and non-IDR regions. The density plot shows that most TFs and most non-TFs have elevated θ_W_ in their IDRs. **d**, Nucleotide diversity θ_W_, calculated for the TF genome and non-TF genome. **e**, Nucleotide diversity π, in TFs (*n =* 1,483; left) and non-TFs (*n =* 10,182; right), with between 10% and 90% disorder content. π was calculated separately for the protein’s IDR and non-IDR regions. The density plot shows that most TFs have increased π in their IDRs. For non-TFs, this bias is small, showing higher π in their IDR regions as well. **f**, Nucleotide diversity π, calculated for the TF genome and non-TF genome. **g**, Tajima’s D in TFs (*n =* 1,483; left) and non-TFs (*n =* 10,182; right) with between 10% and 90% disorder content. Tajima’s D was calculated separately for the protein’s IDR and non-IDR regions. **h,** Tajima’s D calculated for the TF genome and non-TF genome. **i,** EAS-specific LSBL-derived F_ST_ values in TFs (*n =* 1,092; left) and non-TFs (*n =* 5,992; right). There is a significant skew towards higher F_ST_ values in a TF’s IDR region compared to its non-IDR region, which is not observed in non-TFs.

Interestingly, although indels exhibit the same qualitative patterns as SNVs, the differences in the density of TF indels between IDR and non-IDR regions (Binomial exact test, *P < 1*0^−30^; 21.7 vs 17.1 indels/kb, respectively) are substantially smaller compared to non-TFs (Binomial exact test, *P < 1*0^−30^; 22.8 vs 18.4 indels/kb in IDRs and non-IDRs, respectively). Therefore, TF IDRs are less tolerant to indels than non-TF IDRs, suggesting that changes in activation-domain length are deleterious despite their propensity to be disordered.

To more quantitatively assess levels of genetic diversity, we calculated θ_W_ and π in IDRs and non-IDRs across the proteome (Fig. 4c-f). θ_W_ measures the number of segregating sites normalized by sequence length, whereas π measures the average number of pairwise nucleotide differences between chromosomes. Under the standard neutral model, both of these statistics estimate the scaled population mutation rate θ = 4*N_e_*μ, where *N_e_* is the effective population size and μ is the mutation rate per bp per generation. In 74.4% of TFs, θ_W_ was higher in IDRs compared to non-IDRs (Wilcoxon, *P < 1*0^−30^; Fig. 5c). A similar pattern was observed for non-TFs, where 72.4% of genes had elevated θ_W_ values in IDRs (Wilcoxon, *P < 1*0^−30^). The overall difference in θ_W_ between all IDR and all non-IDR regions is larger for TFs compared to non-TFs (Fig. 4d). Similarly, π also varied between ordered and disordered regions of TFs (Fig. 4e-f). Specifically, 65.7% of TFs had higher π in their IDRs compared to their non-IDR regions (Wilcoxon, *P =* 9.3×10^−28^). In contrast, only 49.8% of non-TFs had a higher π in IDRs (Wilcoxon, *P =* 2.9×10^−4^; Fig. 4e-f), likely reflecting the smaller differences in constraint between ordered and disordered regions compared to TFs. Consistent with this hypothesis, the overall difference in π between IDR and non-IDR regions was larger for TFs compared to non-TFs (Fig. 4f).

**Fig. 5.**
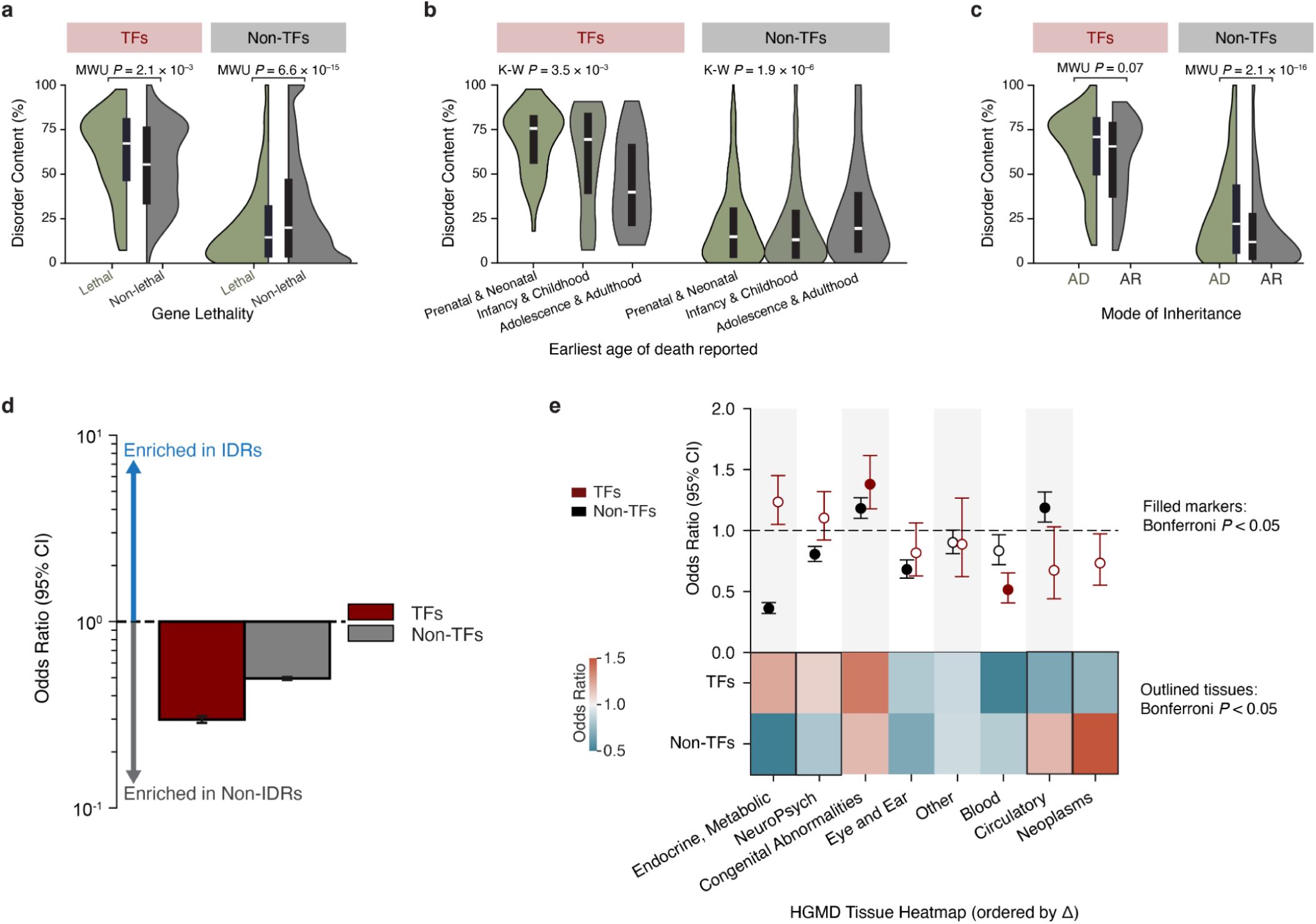
IDRs of TFs and non-TFs exhibit different disease characteristics and burdens. **a**, Disorder content of TFs and non-TFs stratified by OMIM lethality annotation. Lethal TFs show higher disorder content than non-lethal TFs, whereas lethal non-TFs show reduced disorder content relative to non-lethal non-TFs (two-sided Mann–Whitney U tests; TF lethal *n =* 110, TF non-lethal *n =* 1,502; non-TF lethal *n =* 1,529, non-TF non-lethal *n =* 16,595). The black box denotes the IQR, and the white line indicates the median. **b**, Among OMIM-lethal genes, disorder content stratified by the earliest reported age of death. TF disorder content is highest for prenatal/neonatal lethality and decreases for later lethality, while non-TFs show the opposite pattern (Kruskal–Wallis tests; TF *n =* 52, 54, 19; non-TF *n =* 514, 810, 412 for prenatal/neonatal, infancy/childhood, and adolescence/adulthood, respectively). **c**, Disorder content stratified by mode of inheritance (autosomal dominant (AD) versus autosomal recessive (AR)). Disorder content does not differ between AD and AR TFs, whereas AD non-TFs show higher disorder content than AR non-TFs (two-sided Mann–Whitney U tests; TF AD *n =* 113, TF AR *n =* 69; non-TF AD *n =* 621, non-TF AR *n =* 1,593). **d**, Enrichment of HGMD disease variants in IDR versus non-IDR coding sequence for TFs and non-TFs. OR<1 (below the dashed line) indicates relative depletion of disease variants in IDRs (TFs: OR=0.298, 95% CI=0.286–0.310; non-TFs: OR=0.495, 95% CI=0.487–0.504). **e**, Disease-category stratified enrichment of HGMD variants in IDRs within TFs and non-TFs. Points indicate ORs with 95% CIs, and filled markers indicate Bonferroni-significant enrichment. The heatmap summarizes ORs across tissue categories (outlined tiles denote Bonferroni significance), ordered by the ΔOR between TFs and non-TFs.

To better understand the different patterns of π and θ_W_, we calculated Tajima’s D, which is the normalized difference between π and θ_W_. Tajima’s D is expected to be zero under the standard neutral model. Tajima’s D was negative in all comparisons, consistent with an excess of rare variants due to human population growth^58,59^, but IDRs again showed a distinct bias (Fig. 4g-h). For TFs, 59.1% of genes had more negative values in IDRs compared to non-IDRs. Non-TFs showed bias in the other direction, with 73.8% of genes having elevated D in non-IDRs compared to non-IDRs. Across the genome, IDR regions showed more negative D values than non-IDR regions for both TFs and non-TFs (Fig. 4h). We confirmed that these trends were similar using variants from the 1000 Genomes Project^60^ (Extended Data Fig. 7a-h). Together, these findings indicate that IDRs have a higher density of rare variants than non-IDRs, consistent with their increased mutational tolerance, and that these trends were stronger in TFs than in non-TFs.

Finally, we compared patterns of population structure between IDRs and non-IDRs, which may provide insights into how selection shapes variation in these regions. To quantify population structure, we calculated locus-specific branch lengths (LSBL)^61^ in African (AFR), European (EUR), and East Asian (EAS) individuals from the 1000 Genomes Project^60^. LSBL is a simple function of pairwise F_ST_ values among the three populations, yielding a branch-specific F_ST_ value for each population, with higher values indicating greater allele frequency divergence. In the EAS population, we compared per-gene F_ST_ values between IDR and non-IDRs (Δ*F_ST_* = *F_ST, IDR_* − *F_ST, Non−IDR_*). We restricted this analysis to proteins with at least 20% disorder and no more than 80% disorder. For non-TFs, ΔF_ST_ was positive in 49.6% of genes, indicating the magnitude of structure was approximately equal among structured and non-structured regions (Wilcoxon, *P >* 0.05; Fig. 4i). In contrast, ΔF_ST_ was skewed towards positive values for 55.2% of TFs, showing that TF IDRs are enriched for structure (Wilcoxon, *P =* 2.6×10^−4^; Fig. 4i). This bias was also seen in AFR and EUR populations (Extended Data Fig. 7i-j). The most parsimonious explanation for these results is that differences in the intensity of purifying selection between IDR and non-IDR regions is larger for TFs compared to non-TFs (i.e., relative to structured domains, TF IDRs experience more relaxation of functional constraint than non-TF IDRs).

### Disorder content shapes the disease landscape of TFs and non-TFs

We investigated whether the disorder content of TFs and non-TFs was related to disease characteristics. We first analyzed genes associated with lethal phenotypes (see Methods). Lethal TFs had a significantly higher disorder content than non-lethal TFs (Mann-Whitney U test, *P =* 2.1×10^−3^; Fig. 5a), consistent with the observation that highly disordered TFs are enriched for core biological functions (Fig. 2g). In contrast, non-TFs showed the opposite trend with lethal genes being depleted for disorder compared to non-lethal genes (Mann-Whitney U test, *P =* 6.6×10^−15^; Fig. 5a). Among lethal TFs, the median disorder content was significantly higher for proteins that exhibited prenatal or neonatal lethality compared to adolescent or adult lethality (Kruskal-Wallis: *P =* 3.5×10^−3^; Fig. 5b). Again, non-TFs exhibited the opposite pattern with adolescent or adult lethality associated with the highest disorder (Kruskal-Wallis: *P =* 1.9×10^−6^; Fig. 5b). A contributing factor to these differences may be that developmental TFs are significantly more disordered than non-developmental TFs (Extended Data Fig. 4b). More broadly, these opposing trends are consistent with the view that TF IDRs often encode interaction capacity that can have far-reaching regulatory effects, whereas lethal non-TFs may be enriched for proteins whose essential function depends on structured catalytic or structural domains.

We also found that the mode of inheritance for pathogenic mutations was significantly related to disorder content (Fig. 5c). Specifically, genes with autosomal dominant (AD) pathogenic mutations were more disordered than genes with autosomal recessive (AR) mutations for both TFs and non-TFs (Fig. 5c). Although this difference was only marginally significant for TFs (Mann-Whitney U test, *P* = 0.069; Fig. 5c), it was highly significant for non-TFs (Mann-Whitney U test, *P =* 2.1×10^−16^; Fig. 5c) and when TFs and non-TFs were combined (Mann-Whitney U test, *P =* 9.97×10^−29^). Given that the effect sizes for TFs and non-TFs were nearly identical (Fig. 5c), the less significant results for TFs are a consequence of lower power as they have a ~20-fold lower sample size. We hypothesize that disordered proteins are more likely to be AD because IDRs mediate flexible, multivalent interactions; thus, mutations in IDRs are more likely to be dominant-negative, gain-of-function, haploinsufficient, or disrupt the formation of phase-separated condensatesr^62–64^. In contrast, mutations in proteins with less disorder are more likely to cause loss-of-function effects that do not disrupt the activity of the wild-type allele^65–67^, and thus have an AR mode of inheritance.

We next examined whether causal disease variants were more likely to occur in ordered or disordered regions of proteins. High-confidence pathogenic SNVs from the Human Gene Mutation Database (HGMD)^68^ were enriched in ordered regions, consistent with their stronger structural constraints (Fig. 5d). However, this enrichment was weaker for TFs than for non-TFs, indicating that, relative to other proteins, pathogenic SNVs in TFs are more evenly distributed between IDRs and non-IDRs. Thus, TF IDRs carry a higher burden of pathogenic variants than IDRs in the rest of the proteome. This observation suggests that pathogenic variants in TF IDRs are more consequential than previously appreciated, potentially because disruption of IDR-mediated interactions or condensate assembly can have broad regulatory consequences.

Finally, we asked whether the balance of IDR versus non-IDR pathogenic variants differs by disease type (Fig. 5e). Grouping HGMD variants by annotated tissue or disease category, we calculated an odds ratio that quantified the odds of a pathogenic mutation occurring in an IDR versus the odds it occurs in a non-IDR for TFs and non-TFs. We found four disease categories that had significantly different odds ratios between TFs and non-TFs (endocrine/metabolic, neuropsychiatric, blood, and neoplasms; Fig. 5e). Neuropsychiatric phenotypes showed a relative shift towards pathogenic mutations occurring in IDRs in TFs compared to non-TFs, whereas endocrine or metabolic and blood disorders showed stronger enrichment of pathogenic variants in non-IDR regions, particularly for non-TF genes. These tissue-specific patterns suggest that the contribution of TF IDRs to disease is not uniform across phenotypes, and that disruption of TF IDRs is especially relevant for particular disorders.

## Discussion

TFs are fundamentally important proteins that regulate the timing, cell-type specificity, and magnitude of gene expression^1,31^. Through their ability to integrate diverse regulatory inputs and coordinate complex gene regulatory networks, TFs influence nearly all biological processes, and many diseases result from aberrant gene expression^31^. Despite decades of work revealing how TFs bind DNA, recruit cofactors, and modulate chromatin and transcriptional output, our understanding of the mechanistic principles underlying TF function remains incomplete. In particular, accumulating evidence suggests that gene regulation is often organized through the assembly of transcriptional condensates, dynamic, multicomponent hubs that concentrate TFs, coactivators, and the transcriptional machinery at regulatory elements^14,15^. A defining feature of TFs that enables such assemblies is the presence of IDRs, which mediate multivalent interactions and are thought to play a central role in condensate formation and function. These observations raise the possibility that TF IDRs are subject to distinct functional and evolutionary constraints relative to disordered regions in other proteins. Motivated by this hypothesis, we comprehensively compared the properties of IDRs in TFs and non-TFs, finding that distinct evolutionary pressures and functional demands shape TF disorder.

In contrast to most proteins that evolve towards greater order over time as they acquire energetically favorable secondary structures, we find that TFs systematically deviate from this paradigm. Strikingly, TFs have evolved towards greater disorder. We hypothesize that TFs are selected to maintain or expand their disorder content as their GRNs become increasingly complex, context-specific, and entrenched over time. Ancient TFs are more deeply integrated into regulatory pathways, and selection may favor IDRs that enable multivalency (cofactor recruitment) and more tissue-specific regulatory function. Additionally, TF IDRs can evolve under constraints that deprioritize fold stability in favor of increasing disorder content, thereby conferring a more complex cofactor interaction grammar, such as motif spacing, charge patterning, and the density of aromatic residues. In short, TFs, like most of the proteome, are disordered but different in their functional characteristics and evolution, so proteome-wide generalizations about disorder should be interpreted cautiously. These findings also suggest that IDR-mediated transcriptional condensates are potential contributors to regulatory innovation and human phenotypic evolution.

Across interaction and regulatory networks, TF IDRs are uniquely correlated with regulatory scope and complexity. For example, the disorder content of TFs, but not non-TFs, scales with the size of the physical PPI network, consistent with increased interaction capacity for cofactor recruitment during transcription. Notably, this trend is almost entirely driven by the largest family of TFs, C2H2 zinc finger TFs (Spearman’s *ρ =* 0.55, *P <* 10^−30^; Extended Data Fig. 8a-i). When using disorder content as the total number of disordered residues (rather than the percentage), the PPI correlation was largely, but not entirely, driven by C2H2 zinc-finger TFs (Spearman’s *ρ =* 0.40, *P =* 5.7×10^−30^; Extended Data Fig. 8b). Other results, including tissue-specific expression, inter-individual expression variance, and number of tissues with an eQTL, were not driven by a single TF family (Extended Data Fig. 8c-i).

Furthermore, the disorder content of TFs scales with increasing regulatory complexity, as demonstrated by positive correlations with the size of the functional association PPI network, the number of target genes, and the number of tissues in which a TF governs a tissue-specific expression pattern. Interestingly, these trends emerge from IDRs rather than from structured DBDs, suggesting that IDR differences, rather than DBD differences, better capture variation between TF GRNs.

Additionally, our data suggests an interaction between IDRs and tighter dosage control of TFs themselves. Disorder content in TFs is associated with tighter transcript abundance and isoform regulation, reflected in reduced inter-individual expression and splicing variability and a depletion of fine-mapped eQTL and sQTL signals. One explanation is that IDR-mediated multivalency increases a TF’s regulatory influence, so fluctuations in dosage or isoform composition propagate across more targets and contexts, increasing fitness costs and selecting for regulatory mechanisms that stabilize TF abundance.

Genetic variation in IDRs also differs between TFs and non-TFs. IDRs show higher nucleotide diversity (π and θ_w_) compared to non-IDRs, with a larger contrast for TFs rather than non-TFs. Similarly, population differentiation is elevated in IDRs compared to non-IDRs in TFs, but not in non-TFs. Extending to human disease, TFs associated with lethal Mendelian phenotypes tend to have higher disorder and earlier developmental lethality, suggesting that coding variation in TF IDRs can contribute to severe phenotypes even though the overall pathogenic burden remains enriched in non-IDRs. Current variant-effect predictors rely heavily on sequence conservation, which is often weak in IDRs even when these regions are functionally constrained. Thus, genetic variation and the pathogenic burden of TF IDRs are structured by TF regulatory context, such as network connectivity and tolerated expression variation. Therefore, incorporating these gene- and network-level features alongside local sequence properties may improve variant-impact prediction in IDRs beyond conservation alone.

There are several limitations of this work. For instance, although the methods we used to predict disorder have been shown to perform well^32,35,69^, future studies will ideally leverage experimentally validated IDRs. Furthermore, the binary classification of a protein segment as being disordered or ordered does not reflect the complexity of IDRs, which can undergo context-dependent order-to-disorder transitions and vice versa. Finally, isoform-aware analyses are needed to resolve cell-type-specific regulatory activity and to avoid conflating protein- and transcript-level effects. Future work integrating improved catalogs of structural variation, isoform-resolved annotations, and analyses of conservation of biochemical grammar would strengthen mechanistic inference.

In summary, TF IDRs have unique functional, phenotypic, and evolutionary characteristics, and our results provide novel insights into how protein disorder shapes the evolution of complex gene regulatory systems. More broadly, mapping how variation in TF IDRs uniquely rewires regulatory interactions provides a foundation for interpreting mutations in these regions and understanding how regulatory dysfunction arises in human disease.

## Methods

### Obtaining and Curating the Human Proteome

The UniProt reference human proteome was downloaded from the UniProt Knowledgebase. This dataset (Proteome ID: UP000005640; Taxonomy: 9606; Organism: Homo sapiens; Release: 2025_04; Date: 15-Oct-2025) initially comprised 20,659 proteins with canonical sequences. Protein annotations were obtained from the UniProt API^70^. We applied a series of filters to retain high-confidence protein-coding genes. Specifically, entries not included in the manually curated Swiss-Prot (SP) database were excluded. Additional exclusions were made for proteins lacking gene name annotations, those shorter than 20 amino acids, and proteins with the lowest UniProt protein existence score (score=5). Post-filtering, a total of 19,736 proteins were retained.

To identify the subset of proteins with genomic coordinates from the GRCh38/hg38 reference, we linked UniProt accession IDs to CDS-level annotation via the Proteins REST API^71^ (https://www.ebi.ac.uk/proteins/api/coordinates). Proteins without genomic coordinates and those that did not map to chromosomes 1–22, X, and Y, were discarded. This process yielded a set of 19,226 proteins. To confirm the accuracy of these coordinates, we retrieved the GRCh38 sequences corresponding to each protein’s CDS coordinates using the Ensembl REST API (https://rest.ensembl.org/sequence/region/human), resulting in a final set of 18,899 proteins (18,077 autosomal, 785 X-chromosomal, and 37 Y-chromosomal proteins). To connect UniProt entries to entries in other genomic databases, UniProt ID-to-Ensembl Gene Stable ID mappings were used.

### Annotation of Transcription Factors

A total of 1,639 TFs and their corresponding DNA-binding domain (DBD) motifs were identified from the manually curated Human TFs catalog (2018, v1.01; http://humantfs.ccbr.utoronto.ca)^31^. An intersection of the 1,639 TFs with the aforementioned 19,736 UniProt proteins, using Ensembl IDs, resulted in a refined set of 1,613 TFs for subsequent analyses. The set of pioneer TFs was obtained from a previously published list^48^. Our TF set included 32 out of 34 pioneer TFs listed in this resource (*PAX7, PBX1, GRHL2, GATA4, FOXO1, GATA5, SPI1, GRHL1, CEBPB, SOX9, SOX2, FOXA1, GATA1, GATA2, FOXM1, RUNX3, FLI1, FOXD3, TFAP2C, GATA3, GATA6, PBX3, POU5F1, ASCL1, TP63, RUNX1, MYB, FOXE1, TP53, OTX2, FOXA2, KLF4*).

### Identifying IDRs

A consensus disorder prediction tool, MetaPredict (v3.1), was used to identify IDRs, which has been shown to be highly accurate in benchmarking studies^34^. The UniProt FASTA file of primary sequences was input into the MetaPredict Colab notebook (https://colab.research.google.com/github/holehouse-lab/ALBATROSS-colab/blob/main/idrome_constructor/idrome_constructor.ipynb), which assigned disorder scores between 0 and 1 for each residue and used the default score threshold of ≥ 0.5 to classify residues as disordered. In defining IDRs, we used a minimum size threshold of 20 contiguous amino acids to focus on functionally relevant segments and avoid short linker regions. These criteria yielded 28,565 IDR segments.

We also called proteome-wide IDRs on the same UniProt FASTA file using the AIUPred software with the default settings (--analysis-type disorder --no-smoothing False). We selected AIUPred because it exhibits high accuracy in independent benchmarking studies^34^ and is methodologically distinct from MetaPredict. Briefly, AIUPred uses a deep learning algorithm trained on an explicit biophysical model to identify IDRs, whereas MetaPredict obtains consensus calls by integrating many disorder prediction methods. In defining IDRs, we once again used a score threshold of ≥ 0.5 and a minimum size threshold of 20 contiguous amino acids. These criteria yielded 28,991 IDR segments.

### Gene age estimates

We obtained gene age estimations from GenOrigin^39^, a catalog of gene ages inferred across 565 species. Out of the 19,736 proteins described above, 18,123 had TimeTree^72^ age estimates and Ensembl IDs that mapped to a UniProt accession number. Although gene intervals ranged from ‘0–7 Mya’ to ‘>4290 Mya’, we aggregated them into four age bins (0–100, 100–500, 500–1000, >1000 Mya) to mitigate potential estimation errors. To confirm the robustness of our results, we used another human gene age catalog^43^ that used GenEra^73^, which accounts for biases introduced by homology detection failure^40^. We mapped Ensembl protein IDs to UniProt accession numbers and filtered out proteins with incomplete phylostratum information and those with phylostrata more recent than Eutheria.

Furthermore, we also repeated analyses by stratifying the data by conservation, protein length, GC content, expression levels, and tissue specificity. We defined conservation using PhyloP 100-way scores downloaded from UCSC (May 2015 version). We intersected this with the aforementioned set of 18,899 proteins with precise genomic coordinates and calculated the per-gene mean phyloP score. These scores were stratified into tertiles in order to test the robustness of the disorder-evolution relationship in TFs and non-TFs. Expression levels and tissue specificity were obtained from GTEx v10 bulk tissue expression data. Specifically, we first obtained the median gene-level TPM by tissue (GTEx_Analysis_v10_RNASeQCv2.4.2_gene_median_tpm.gct.gz). Next, to bin proteins by gene expression level, we calculated the median TPM across all tissues and stratified these values into tertiles. The tissue specificity index, *τ*, was calculated across all tissues according to the original method^74^.

We modeled gene age (from GenOrigin estimates) using ordinary least squares (OLS) regression to test whether the association between gene age and disorder content differs between TFs and non-TFs. TF status was encoded as a binary indicator variable 𝟙[*TF*], (1 for TFs, 0 otherwise). Disorder content *D* (percent disordered residues; 0-100) was mean-centered (*D_C_* = *D* − *D̄*). The interaction term of these two variables was included. The model also included covariates *X*_k_ for protein length, GC content, mean expression level, and tissue specificity (*τ*). We modeled gene age as follows:

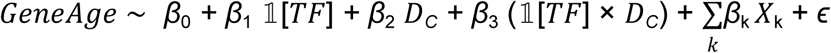

### GO Analysis

Gene Ontology (GO)^75^ enrichment of biological process (BP) terms was performed using GOATOOLS^76^ with reference annotations from the GO Association File (GAF) (https://geneontology.org/docs/download-go-annotations, release from 2025-07-22). Developmental TFs and non-TFs were labeled based on GO term annotation for “developmental process” (GO:0032502). The full GO term ontology OBO file was obtained from the Gene Ontology Knowledgebase (https://geneontology.org/docs/download-ontology, downloaded 2025-07-25). A custom GO slim set was constructed by subsetting to terms at Levels 3 and 5 to avoid overly broad and overly narrow terms. The default GOATOOLS settings were used, with Fisher’s exact test and multiple-testing correction by the Benjamini–Hochberg method to control the false discovery rate. The background set used was the full TF list (*n* = 1,005).

### PPI network analysis

Protein–protein interaction (PPI) data were obtained from the STRING^77^ database (v12.0) after filtering to Homo sapiens-specific data and removing redundant interaction pairs (ex. AB-BA). Analyses were performed using both the full network and the physical subnetwork. To assess robustness, we repeated analyses using multiple interaction confidence thresholds based on the interaction score, which ranges from 0 to 1000. We assessed networks after thresholding to interactions with scores greater than 150, 400, and 700, corresponding to low, medium, and high confidence, respectively. After intersecting this catalog with our protein set using STRING protein IDs and subsetting to proteins with at least 1 interaction, 19,058 proteins were used for further analysis.

We also assessed PPI using the Human Protein Atlas (HPA)^51^ Interaction resource, which integrates data from multiple databases to construct a consensus network of interactions present in at least two datasets. We subsetted to proteins with at least 1 interaction in the catalog and removed proteins with a reliability score of “Uncertain”, resulting in a set of 15,095 proteins. Lastly, we used physical protein interaction counts from BioGRID^52^, resulting in 18,325 proteins with at least 1 interaction.

### GRN analysis

TF–target gene regulatory interactions were obtained from GRNdb^54^, subsetted to TF-gene interactions with “High” confidence. We used regulons predicted from both GTEx (27 tissues) and TCGA (43 tissues) databases. To obtain the total number of targeted genes per TF, we summed the number of unique genes across all tissue regulons. Tissue breadth was defined as the number of distinct tissues in which a TF was connected to at least one target gene.

### Expression Analysis

We obtained gene expression specificity information from the HPA^51^ (version 25.0) and removed proteins with a reliability score of “Uncertain”. For tissue specificity, we used HPA RNA tissue specificity categories (tissue enriched, group enriched, tissue enhanced, low tissue specificity, not detected) and binarized genes as tissue-specific (enriched/group enriched/tissue enhanced) versus broad (low specificity/not detected). For cell-type specificity, we used HPA single cell type specificity categories and similarly binarized to cell-type-specific versus broad.

Inter-individual gene expression variance data were obtained from a previously published dataset^78^, which quantified inter-individual expression variance data from 24 GTEx tissues, 11 TCGA (The Cancer Genome Atlas) tissues, and 22 other individual studies. We intersected genes using Ensembl gene IDs and compared the top 50% “High” and “Low” variance TFs and non-TFs. To show the robustness of the variance-disorder relationship in TFs and non-TFs, we also used the per-study mean gene expression variance to calculate Spearman correlation within each of the 58 tissues and used Bonferroni *P* value correction.

### QTL analysis

We obtained fine-mapped expression QTL (eQTL) and splicing QTL (sQTL) data from GTEx^53^ v10 (November 2024) found across 50 GTEx tissues. Although QTL credible set (CS) overlap was extremely rare (occurring in only 17 instances), we combined any overlapping CSs by at least one variant to prevent redundancy. 17,085 out of 19,736 genes intersected in GTEx genes, based on HGNC ID and ENSG. For each gene, we counted the number of tissues where it was an eGene (linked to at least 1 CS). For each gene, we also counted the number of distinct CSs per tissue to calculate the per-tissue Spearman correlations between eGene disorder and the number of independent CSs. Bonferroni *P* value correction was used. As a summary measure, for each eGene, we identified the tissue with the most credible sets and correlated this maximum CS number with the eGene’s disorder content.

### Translational Efficiency

Translational efficiency (TE) data were obtained from a previously published resource^56^, which predicted TE for 10,037 transcripts across 78 studies, trained on 1,076 human ribosome profiling datasets. We intersected this gene catalog using gene symbols, resulting in 9,887 genes. First, we calculated the per-study Spearman correlation between predicted TE and disorder content, and we used Bonferroni *P* value correction. We then used the mean predicted TE across all samples to summarize the correlation between TE and disorder content.

### Human Population Genetics Analyses

Human genetic variation data were obtained from gnomAD v4.1.0 and the 1000 Genomes Project^60^ (1kGP; Phase 3, accessed March 12, 2019). We used only high-quality variants (FILTER=PASS). From gnomAD, we only used polymorphisms within the non-Finnish European (NFE) subpopulation (AC_nfe>0), in order to minimize population structure effects. From the 1KGP, we used the phased biallelic SNP+INDEL call set for unrelated samples (*n*= 5,248) based on the GRCh38 assembly. To calculate the SNV and indel rates per kilobase, we annotated VCFs by TF status and IDR status (TF IDRs, Non-TF IDRs, TF non-IDRs, and non-TF non-IDRs). We summed variant counts genome-wide within each of the four regions and normalized by the corresponding total genomic region.

To calculate per-base nucleotide diversity metrics, we subsetted to biallelic SNVs. We used the following method to calculate Watterson’s theta (θ*_W_*), where *S* is the number of segregating sites, and *N* is the maximum number of chromosomes in the sample:

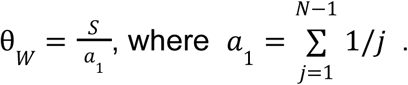

Unlike θ*_W_*, nucleotide diversity (π) is influenced by allele frequencies, and since the number of chromosomes sampled is not consistent across all genomic loci, we weighted the contribution of a single nucleotide by the single-locus number of chromosomes sampled. We obtained π and the standard error using the following formula, where *p_i_* is the frequency of the major allele at locus *_i_*, *q_i_* is the frequency of the minor allele at locus *_i_*, *n_i_* is the total count of chromosomes at locus *_i_*, and L is the total region length:

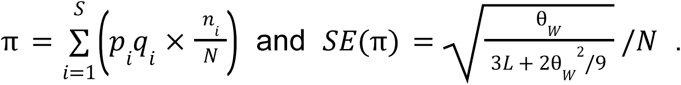

To calculate the per-base π and θ*_W_*, we divided these values by *L*.

We calculated Tajima’s D as follows:

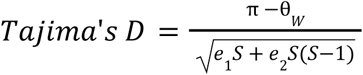

with constants *e*_1_ and *e*_2_ defined as in the original method^79^.

To obtain per-gene metrics and compare the properties of the IDR-coding sequence and the non-IDR-coding sequence within the same gene, we subsetted to proteins with a disorder content between 10–90%. For gnomAD, we used proteins with at least 10 SNVs in each of the protein’s IDR and non-IDR regions; for 1KGP, we used a threshold of 1 SNV.

Using biallelic SNVs from the 1KGP, we estimated unbiased pairwise *FST* values between European (EUR), East Asian (EAS), and African (AFR) populations as described by Weir and Cockerham^80^, using scikit-allel^81^. We used genes with disorder content between 20–80% and compared per-gene IDR *FST* values and per-gene non-IDR *FST* values for each population pair. We used these pairwise *FST* values to estimate locus-specific branch lengths (LSBLs) for each population as previously described^61^. For example, the LSBL of the EAS population is:

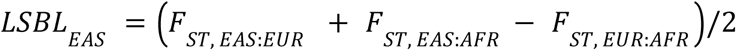

### Functional Variation and Disease Analyses

To obtain genes associated with lethality, we also used the Lethal Phenotypes Portal^82^ (updated November 2024), a catalog of OMIM genes related to lethality. We grouped the Human Phenotype Ontology (HPO)^83^ lethality category annotations to assign the following developmental categories: prenatal/neonatal (L1/L2), infancy/childhood (L3/L4), and adolescence/adulthood (L5/L6). We combined mode-of-inheritance (MOI) annotations from these lethal genes with previously published MOI catalogs^84,85^. We used only autosomal dominant (AD) and autosomal recessive (AR) genes and excluded those that were categorized as both AD and AR.

As an orthogonal source of disease-associated mutations, we also used the Human Gene Mutation Database (HGMD) Professional^68^ (accessed May 2025). We filtered for SNVs with the highest confidence disease-causing mutation associations (tag=DM). To analyze disease-type biases, we used HGMD disease variants linked to ICD-10^86^ disease codes in the HGMD phenbase. ICD-10 chapters were grouped into the following larger disease categories: Neoplasms (C), Blood (D), Endocrine/Metabolic (E), Neuropsychiatric (F, G), Eye/Ear (H), Circulatory (I), Congenital Abnormalities (Q), and Other (J, K, L, M, N).

## Data Availability

The UniProt proteome is available at https://ftp.uniprot.org/pub/databases/uniprot/current_release/knowledgebase/reference_proteo mes/Eukaryota/UP000005640/UP000005640_9606.fasta.gz. UniProt ID mappings are available at https://ftp.uniprot.org/pub/databases/uniprot/current_release/knowledgebase/reference_proteo mes/Eukaryota/UP000005640/UP000005640_9606.idmapping.gz. The TF list is available at https://humantfs.ccbr.utoronto.ca/download/v_1.01/DatabaseExtract_v_1.01.csv. The GenOrigin human gene age catalog can be downloaded at http://chenzxlab.hzau.edu.cn/GenOrigin/#!/download. The GenEra-produced gene age catalog can be downloaded at https://github.com/LotharukpongJS/phylomapr/blob/main/data-raw/Homo_sapiens_gene_ages. tsv. PhyloP scores were obtained from https://hgdownload.cse.ucsc.edu/goldenpath/hg38/phyloP100way/hg38.phyloP100way.bw. The GO association file can be found at https://current.geneontology.org/annotations/goa_human.gaf.gz. The GO OBO file can be found at https://purl.obolibrary.org/obo/go/go-basic.obo. The STRING PPI database is at https://string-db.org/cgi/download. The GRNdb is available at http://www.grndb.com. The Human Protein Atlas (v25.0) is available at https://www.proteinatlas.org/. GTEx v10 QTLs can be downloaded from the GTEx Portal at https://gtexportal.org/home/downloads/adult-gtex/qtl. Translation efficiency data can be found at Zheng et al. (2025) in Supplementary Table 1 at https://doi.org/10.1038/s41587-025-02712-x. VCF files from gnomAD v4.1 were downloaded using Google Cloud Public from https://gnomad.broadinstitute.org/data. 1000 Genomes Project biallelic SNV and indel VCFs (release 2019) were obtained from their FTP site at /vol1/ftp/data_collections/1000_genomes_project/release/20190312_biallelic_SNP_and_INDE L. Lethal OMIM genes data are available at the Lethal Phenotypes Portal https://lethalphenotypes.research.its.qmul.ac.uk/. HGMD**®** Professional data were obtained from https://www.hgmd.cf.ac.uk/ac/index.php.

## Code Availability

Code to reproduce the results is available at https://github.com/AkeyLab/tf_idr_paper.

## Acknowledgements

We would like to thank the members of the Akey laboratory for their helpful comments and feedback on this work, especially Rob Bierman, Cara Weisman, and Aaron Pfennig. We would also like to thank Thouis Ray Jones, Jap Chae, and Jjam Ppong. This work was supported in part by National Institutes of Health grants T32HG003284 (SS) and R01GM110068 (JMA).

## Extended Data Figures

**Extended Data Fig. 1.**
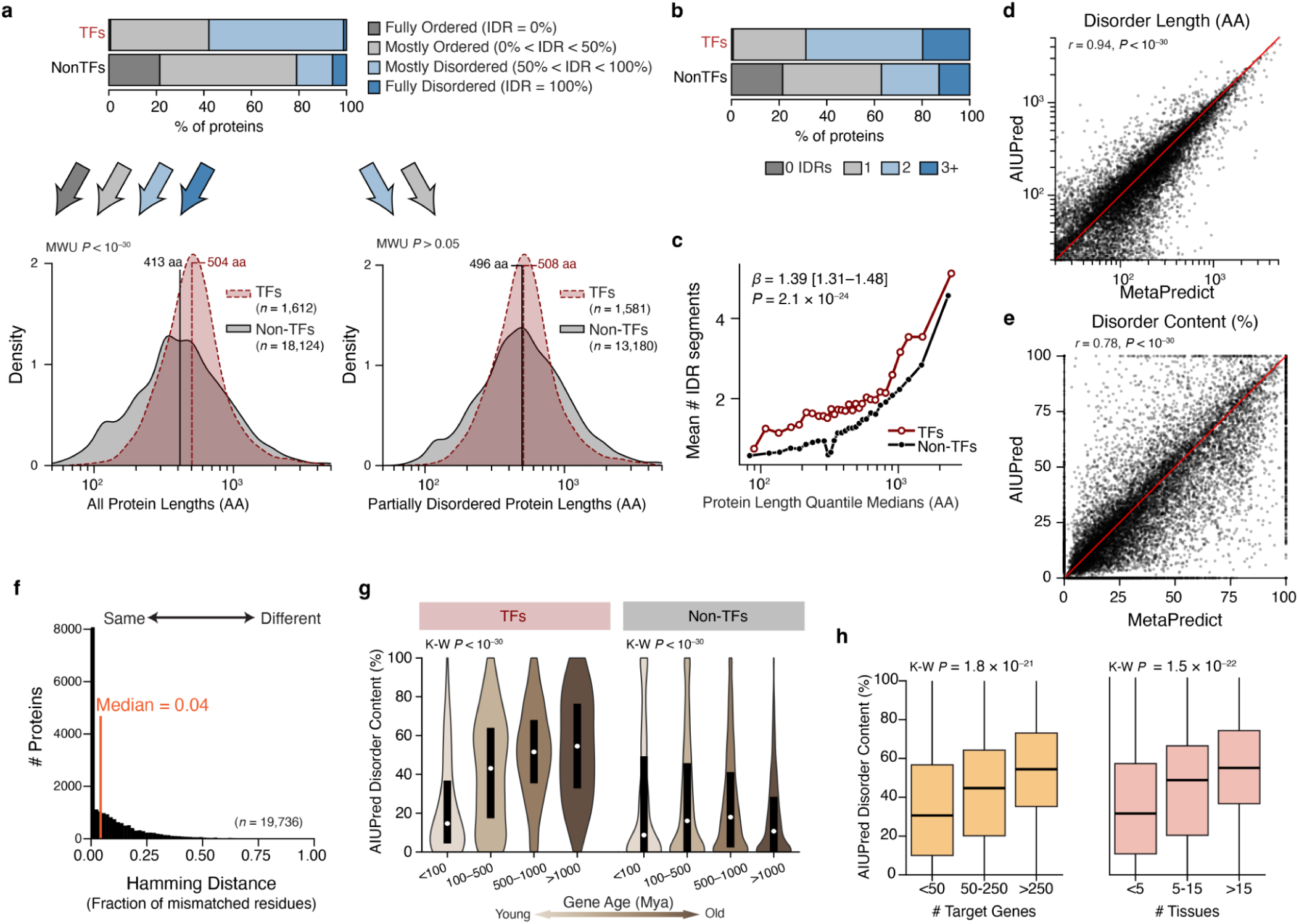
Additional IDR characteristics and robustness across disorder predictors. **a,** Distribution of disorder categories for TFs and non-TFs based on the fraction of residues classified as disordered (fully ordered, 0% IDR; mostly ordered, <50%; mostly disordered, >50%; fully disordered, 100%). Protein-length distributions for all proteins (bottom left) and for partially disordered proteins only (bottom right). Vertical lines denote group medians. *P* values are from two-sided Mann–Whitney U tests. **b**, Distribution of the number of IDR segments per protein (0, 1, 2, or ≥3) in TFs and non-TFs. **c**, Mean number of IDR segments across protein-length quantiles (*x* axis shows quantile medians, aa). Effect estimate for TF status is shown (*β*=1.39 [1.31–1.48], *P =* 2.1×10^−24^; model details in Methods). Across almost all quantiles, TFs have a higher mean number of IDR segments despite having similar protein lengths. **d,e**, Agreement between MetaPredict and AIUPred, a disorder predictor that doesn’t rely on conservation information, but solely energy estimation, for predicted disorder length (aa; **d**) and disorder content (%; **e**) across proteins; red line indicates y=x. **f,** Per-protein Hamming distance between binary residue-level disorder calls from MetaPredict and AIUPred (fraction of mismatched residues; median = 0.04; *n =* 19,736 proteins). **g,** AIUPred disorder content (%) versus gene age (Mya) for TFs and non-TFs. Violins show distributions within age bins; black bars and white lines show the IQR and median, respectively. Kruskal–Wallis tests are shown. **h,** AIUPred disorder content (%) stratified by regulatory complexity among TFs with available annotations, shown by the number of target genes and the number of tissues with regulatory activity. Boxes indicate the median and IQR; whiskers denote 1.5× IQR. Kruskal–Wallis tests are shown.

**Extended Data Fig. 2.**
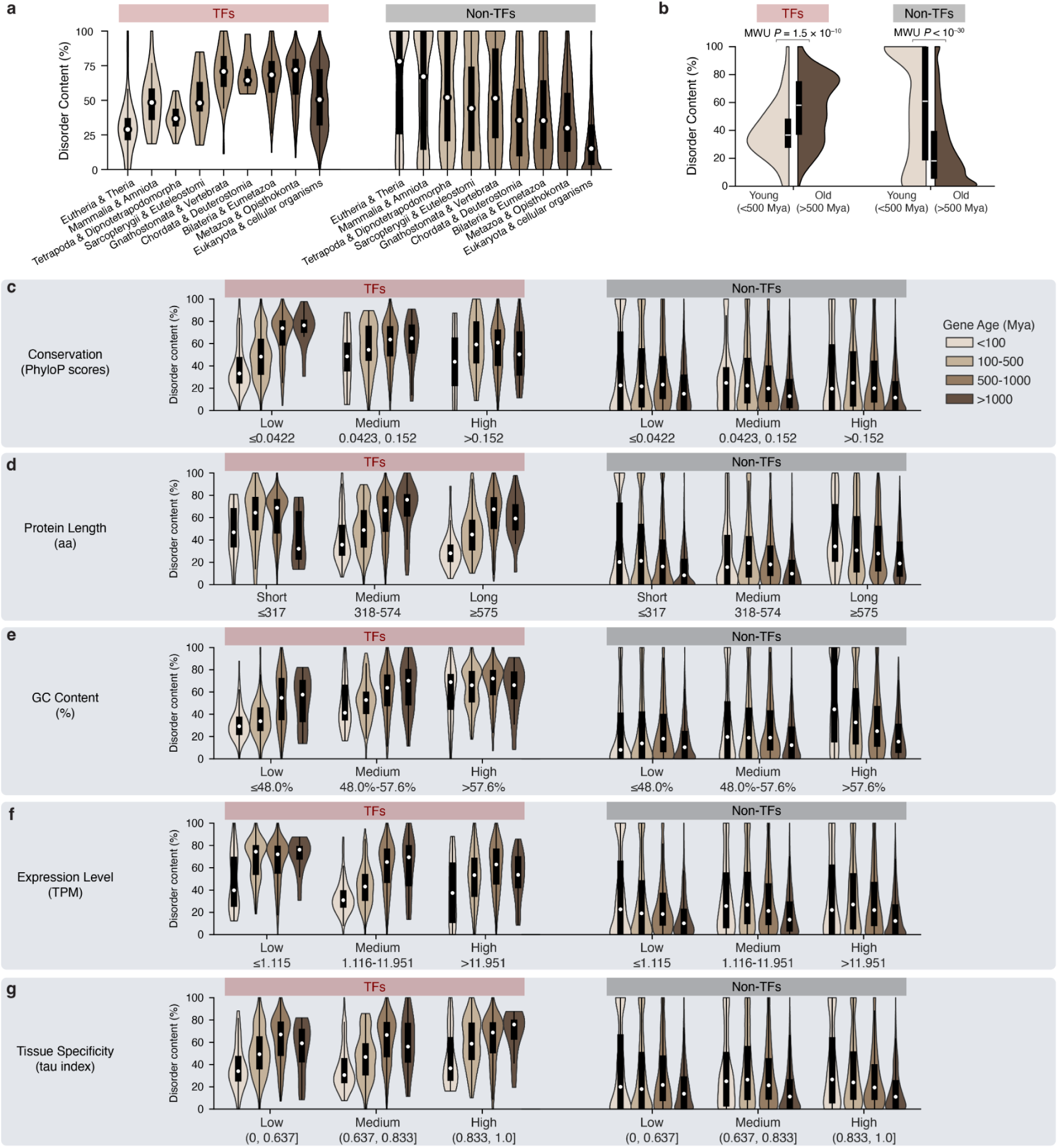
Robustness of the gene age-disorder relationship. **a,** Disorder content in TFs stratified by gene ages estimated by a previous gene catalog^43^ using genEra^73^, a homology-detection failure-aware estimator. Predicted phylostrata, ordered from youngest to oldest clade, are shown separately for TFs (left) and non-TFs (right). **b**, Summary comparison of disorder content for “young” (grouping the youngest 10 phylostrata, Eutheria to Vertebrata, approximately <500 Mya) versus “old” (grouping the oldest 8 phylostrata, Cordata to cellular organisms, >500 Mya) genes within TFs and non-TFs. **c–g,** Stratified analyses of the gene age-disorder relationship, using gene ages from GenOrigin (Fig. 2b), controlling for potential confounders. For TFs (left) and non-TFs (right), their disorder content distribution is shown across gene-age bins within tertiles of **(c)** conservation, **(d)** protein length, **(e)** GC content, **(f)** expression level, and **(g)** tissue specificity.

**Extended Data Fig. 3.**
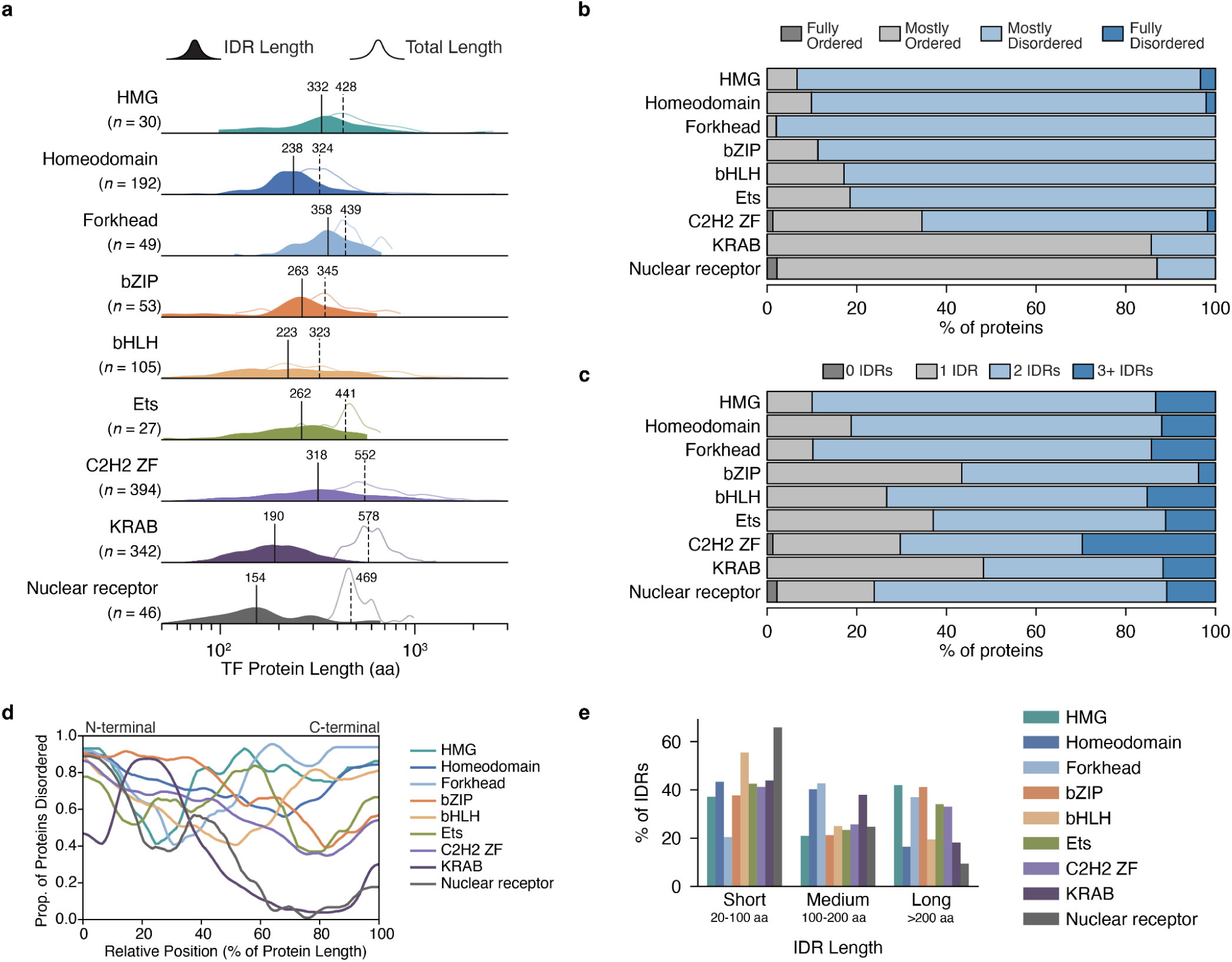
TF IDR features by DNA-binding domain family. **a**, Ridgeline distribution of total protein length (outline) and cumulative IDR length per protein (filled) for major TF DBD families (HMG, Homeodomain, Forkhead, bZIP, bHLH, ETS, C2H2 ZF, KRAB, and Nuclear receptor). This shows that the trends in TF disorder content are not just driven by protein length variation. Vertical lines indicate family medians for IDR length (solid line) and total length (dashed line). **b**, Fraction of proteins in each DBD family classified by disorder content: fully ordered (IDR content=0%), mostly ordered (0%<IDR content<50%), mostly disordered (50%<IDR content<100%), and fully disordered (IDR content=100%). **c**, Distribution of the number of distinct IDR segments per protein (0, 1, 2, or ≥3 segments) within each DBD family. **d**, Positional distribution of disorder along TF sequences by DBD family. Proteins were scaled to relative length (0–100%, N-terminus to C-terminus), and the *y* axis shows the proportion of proteins that are disordered at each position (based on residue-level disorder calls). **e**, IDR lengths by DBD family, shown as the percentage of segments in short (20–100 aa), medium (100–200 aa), and long (>200 aa) categories.

**Extended Data Fig. 4.**
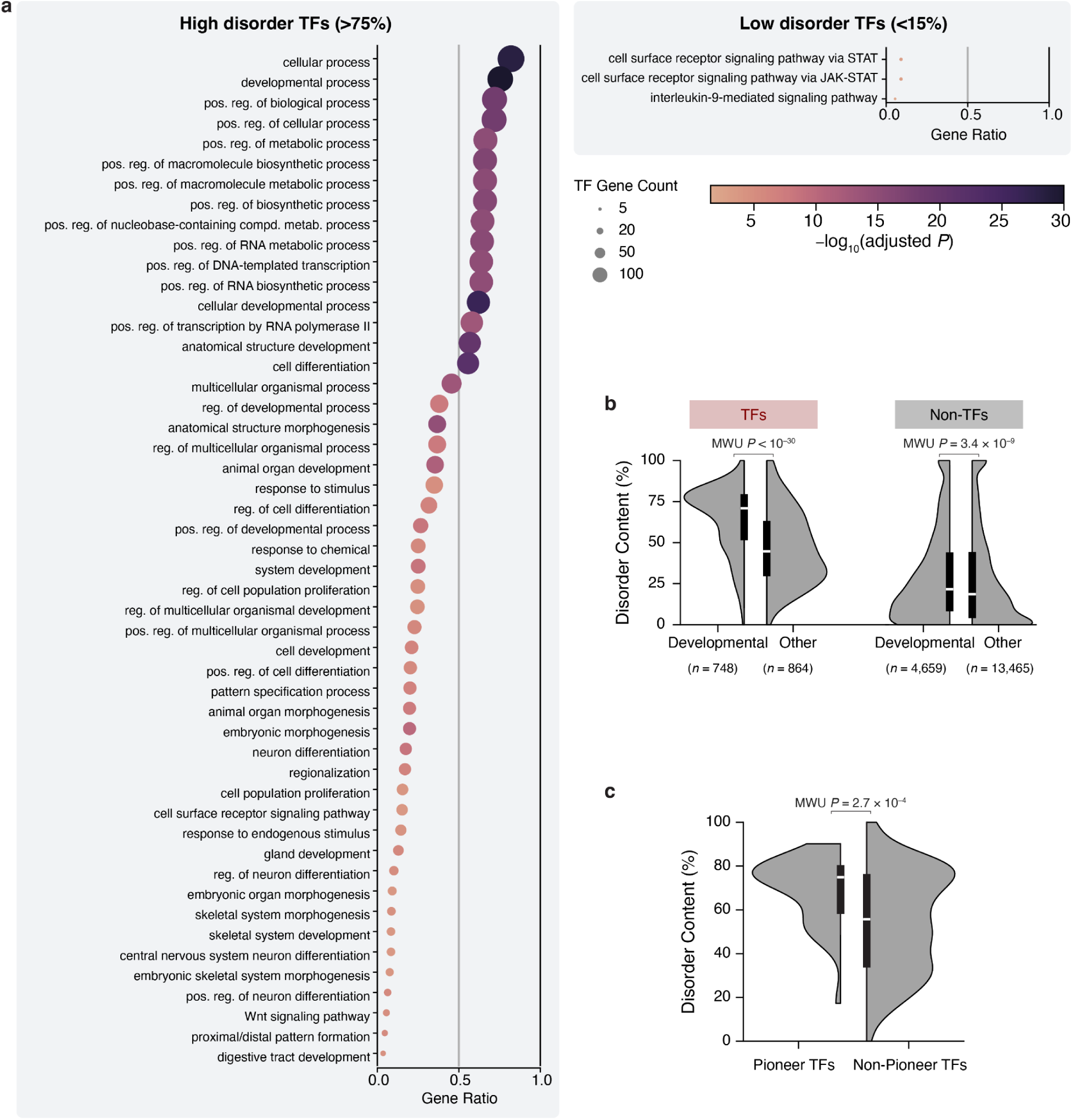
Functional specialization of high- versus low-disorder TFs. **a**, GO Biological Processes enrichment using all GO terms in TFs with high disorder (IDR content>75%; *n =* 351; top) and low disorder (IDR content<15%; *n =* 31; bottom). Dot plots show enriched GO terms (*y* axis). Gene ratio (*x* axis) is the proportion of TFs in the queried set annotated to a given term. The full TF list was used as the background gene set (*n =* 1,008). Dot size indicates the number of TFs annotated to each term, and dot color indicates *−log_10_*(Benjamini–Hochberg adjusted *P* value). **b**, Disorder content (%) for TFs (left) and non-TFs (right) stratified by annotation to “developmental process” (GO:0032502). Black boxes indicate the interquartile range, and the white line indicates the median. The two–sided Mann–Whitney U test *P* value is shown. **c**, Disorder content (%) for pioneer versus non-pioneer TFs. Black boxes indicate the interquartile range, and the white line indicates the median. The two–sided Mann–Whitney U test *P* value is shown. Pioneer TFs were obtained from a previously published dataset^48^.

**Extended Data Fig. 5.**
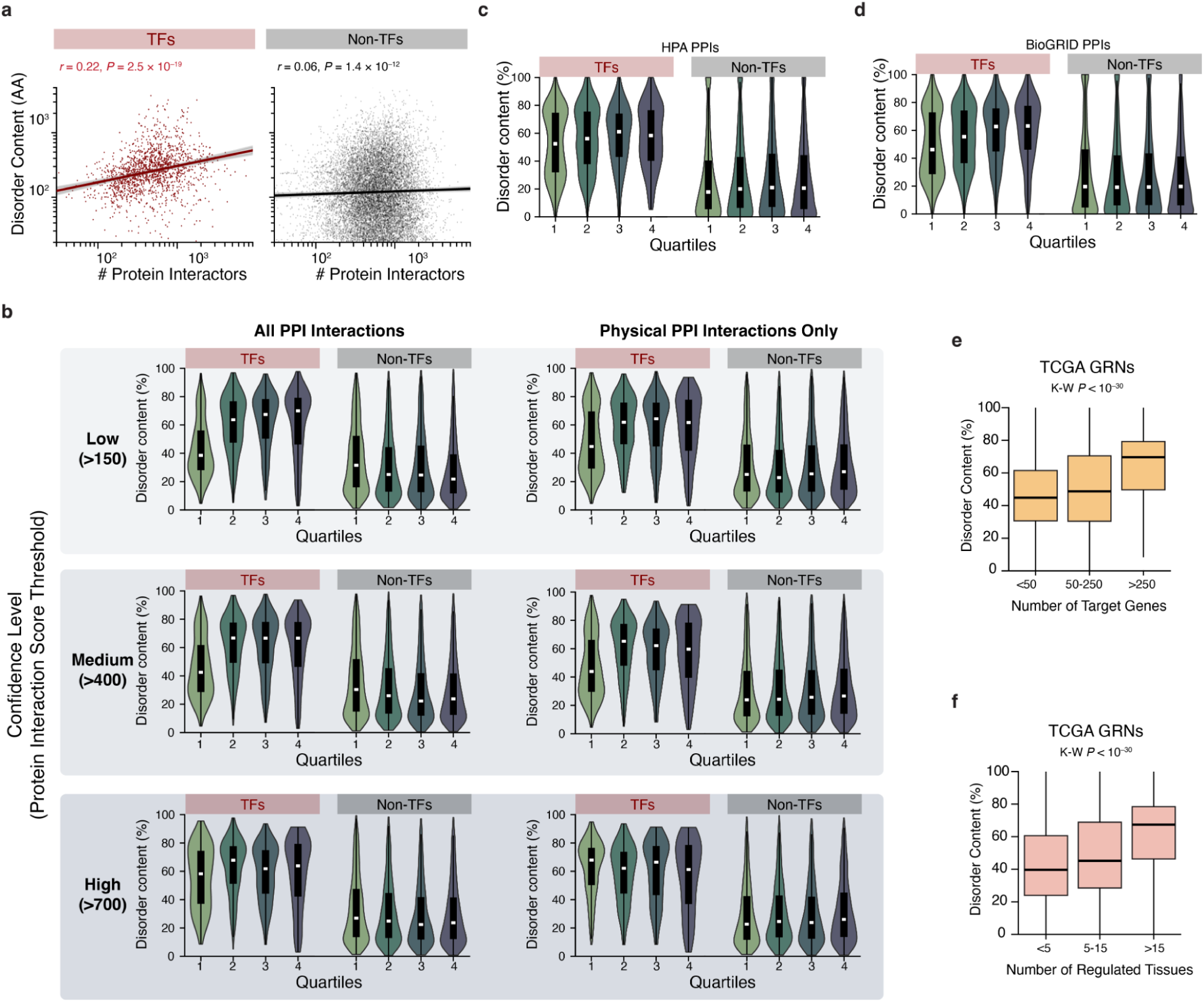
Robustness of the relationship between disorder content and PPI network size. **a,** The PPI-disorder relationship persists when disorder content is measured as the total number of disordered residues per protein rather than as a percentage. We compare TFs (left; red) to non-TFs (right; black). **b,** The PPI-disorder relationship remains when different confidence level thresholds are applied to STRING networks. This shows that the relationship is not purely driven by low-confidence functional links. Disorder content (%) across quartiles of per-protein functional association partner size, shown for TFs and non-TFs, and repeated across STRING confidence thresholds (low>150, medium>400, high>700). Disorder content (%) across quartiles of PPI size constructed using STRING physical (binding) interactions only. **c–d,** The PPI-disorder relationship remains when using different PPI databases. **(c)** Disorder content and quartiles of protein physical interactions estimated from the HPA database. **(d)** Disorder content and quartiles of protein physical interactions estimated from the BioGRID^52^ database. **e,** Disorder content (%) versus the number of target genes in TCGA-derived GRNs. TFs are stratified by the number of inferred target genes per TF (≤50, 51–250, >250). **f,** Disorder content (%) versus regulatory breadth across tissues (as in Fig. 3b) but in TCGA GRNs instead of GTEx GRNs. TFs are stratified by the number of tissues in which a TF is inferred to regulate tissue-specific expression programs (≤5, 6–15, >15).

**Extended Data Fig. 6.**
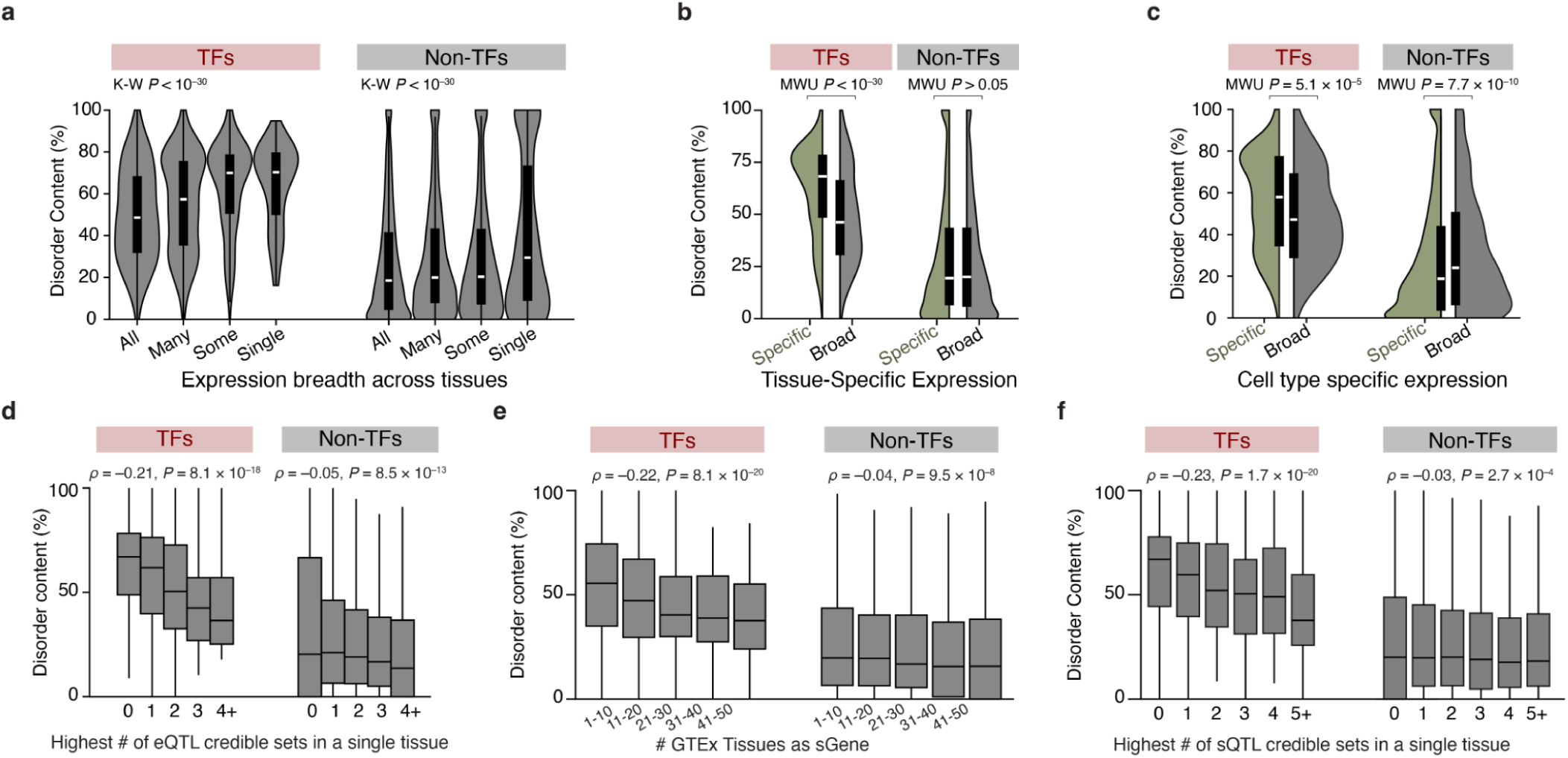
Robustness of associations between disorder content and regulatory breadth across independent datasets and specificity metrics. **a,** Disorder content (%) for TFs and non-TFs stratified by expression breadth across tissues, as annotated by HPA^51^. TFs. *P* values are from the Kruskal–Wallis test. **b,** Disorder content stratified by HPA tissue specificity for TFs (specific TFs *n =* 714; broad TFs *n =* 835) and non-TFs (specific non-TFs *n =* 9,676; broad non-TFs *n =* 7,026). TFs with more uneven expression across tissues show higher disorder content, whereas non-TFs show no significant difference (two-sided Mann–Whitney U tests). **c,** Disorder content (%) for TFs and non-TFs stratified by HPA cell-type specificity (uneven expression across cell types) for TFs. Two-sided Mann–Whitney U tests are shown. **d,** Disorder content (%) stratified by eQTL burden, defined as the maximum number of eQTL credible sets assigned to a gene within any single tissue, shown for TFs (left) and non-TFs (right). **e,** Disorder content (%) stratified by the number of GTEx tissues in which a gene is identified as an sGene (gene regulated by a splicing-QTL), shown for TFs (left) and non-TFs (right). **f,** Disorder content (%) stratified by sQTL (splicing QTL) burden, defined as the maximum number of sQTL credible sets assigned to a gene within any single tissue, shown for TFs (left) and non-TFs (right).

**Extended Data Fig. 7.**
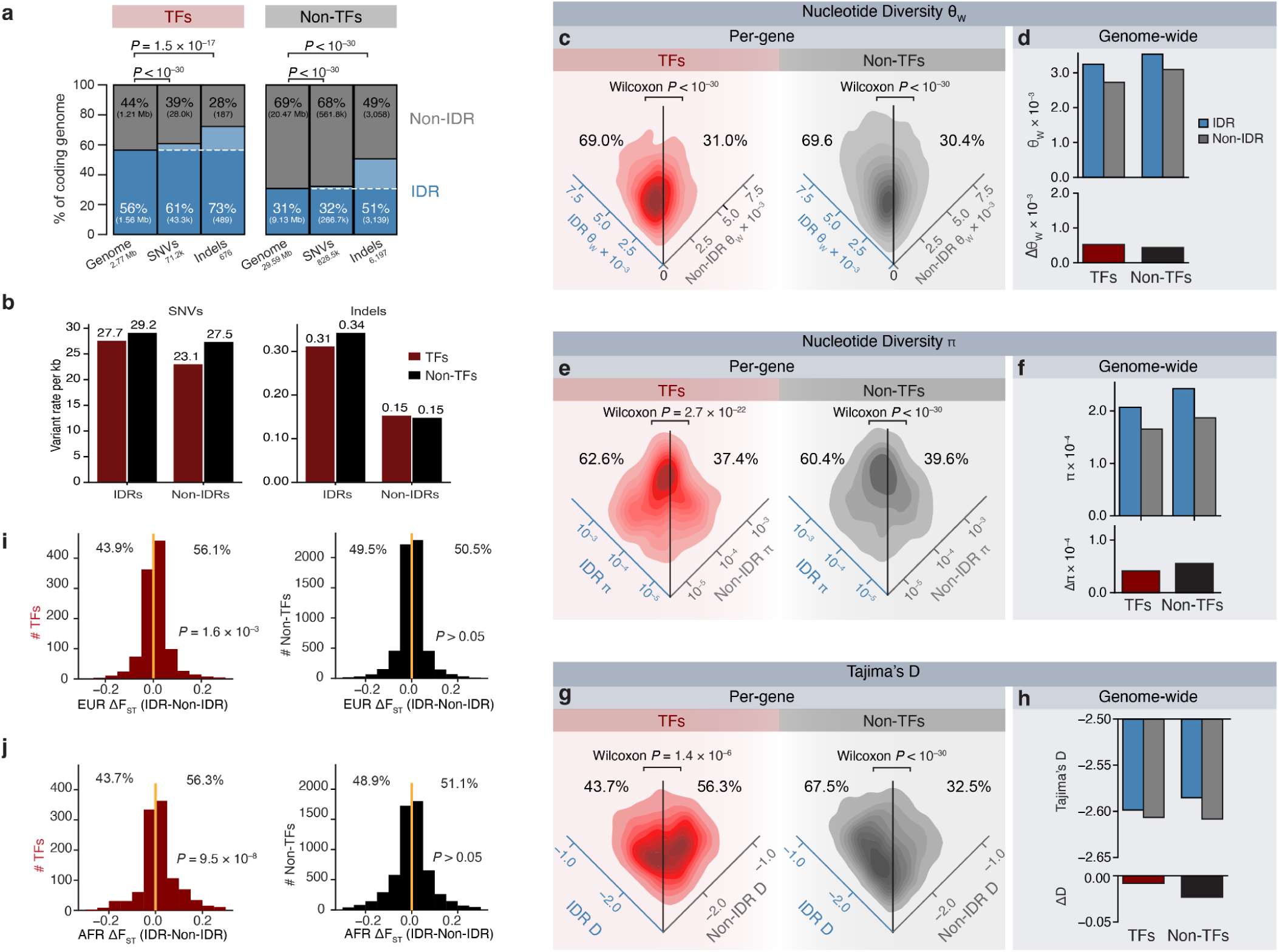
Nucleotide diversity and population differentiation in IDRs versus non-IDRs using 1000 Genomes. **a,** TF IDRs are enriched for SNVs and Indels in the 1000 Genomes Project^60^. Total genomic region, SNV, and indel counts are shown, split between TF Non-IDRs (grey) and IDRs (blue). **b**, SNV and indel rates. **c–d,** Nucleotide diversity (θ_W_) for IDRs and non-IDRs. (**c)** Per-gene θ_W_ estimates for TFs (red) and non-TFs (black) with at least 10% IDR and less than 90% IDR. Density plots compare IDR versus non-IDR values; percentages indicate the fraction of genes with higher diversity in IDRs versus non-IDRs. P values are from two-sided Wilcoxon signed-rank tests comparing paired IDR and non-IDR estimates within genes. (**d)** Genome-wide θ_W_ in IDRs and non-IDRs for TFs and non-TFs (top), with the corresponding genome-wide difference (Δθ_W=_θ_W, IDR_−θ_W,Non-IDR_; bottom). **e–f,** Nucleotide diversity (π) for IDRs and non-IDRs. **(e)** Per-gene π comparisons for TFs and non-TFs with paired Wilcoxon tests and proportions of genes with higher π in IDRs. **(f)** Genome-wide π in IDRs and non-IDRs (top) and the genome-wide difference (Δπ; bottom) for TFs and non-TFs. **g–h,** Tajima’s D in IDRs and non-IDRs. **(g)** Per-gene comparisons of Tajima’s D in IDRs versus non-IDRs for TFs and non-TFs with paired Wilcoxon tests and proportions of genes with higher values in IDRs. **(h)** Genome-wide Tajima’s D in IDRs and non-IDRs (top) and the genome-wide difference (ΔD; bottom) for TFs and non-TFs. **i–j,** Distributions of per-gene differences in population differentiation between IDRs and non-IDRs (ΔF_ST_ = F_ST, IDR_ − F_ST, Non-IDR_) for TFs (*n =* 1,092) and non-TFs (*n =* 5,992) comparing **(i)** Europeans (EUR) and **(j)** Africans (AFR). Vertical lines denote zero difference; percentages and Wilcoxon *P* values indicate the skew towards IDRs or non-IDRs.

**Extended Data Fig. 8.**
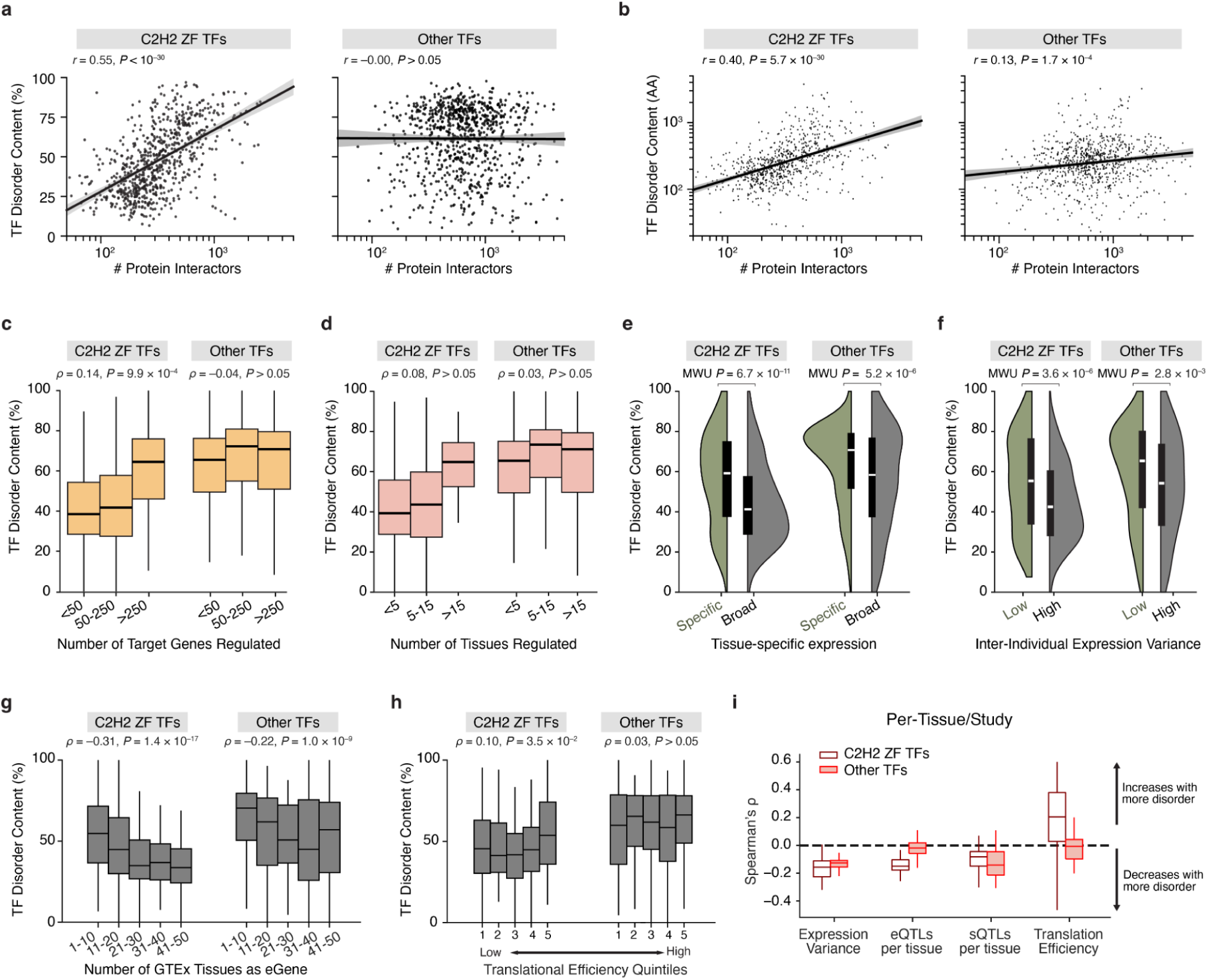
C2H2 zinc finger TFs drive the disorder–network association, while other disorder–functional genomics relationships generalize across TFs. **a,** TF disorder content (%) versus number of physical protein interactors, shown separately for C2H2 zinc finger (C2H2 ZF) TFs and all other TFs; Pearson’s *R* indicated. **b**, Same as in (**a)**, quantifying disorder as the total number of disordered residues per protein (AA) instead of percentage disorder. **c,** TF disorder content (%) stratified by the number of inferred target genes per TF in GTEx-derived gene regulatory networks (GRNs; ≤50, 51–250, >250); Spearman’s *ρ* indicated. **d**, TF disorder content (%) versus regulatory breadth across tissues (as in Fig. 3b) but in TCGA GRNs instead of GTEx GRNs. TFs are stratified by the number of tissues in which a TF is inferred to regulate tissue-specific expression programs (≤5, 6–15, >15). **e,** TF disorder content (%) comparing tissue-specific versus broad expression, shown for C2H2 ZF TFs and other TFs (two-sided Mann–Whitney U test). **f,** TF disorder content (%) comparing genes with low versus high inter-individual expression variance, shown for C2H2 ZF TFs and other TFs (two-sided Mann–Whitney U test). **g,** TF disorder content (%) stratified by the number of GTEx tissues in which a TF is detected as an eGene; Spearman’s *ρ* indicated. **h,** TF disorder content (%) stratified by translational efficiency quintiles (low to high); Spearman’s ρ and two-sided P values indicated. **i,** Per-tissue/per-study associations between TF disorder content and molecular phenotypes, summarized as distributions of Spearman’s ρ across tissues/studies for C2H2 ZF TFs and other TFs; dashed line indicates no association. Box plots show median, interquartile range, and 1.5× IQR whiskers; violins show kernel density with embedded box plots.

## Notes

### Competing Interest Statement

The authors have declared no competing interest.

### Summary of Updates

Figures 3 and 4 have been combined

